# Spatial attention enhances cortical tracking of quasi-rhythmic visual stimuli

**DOI:** 10.1101/689711

**Authors:** D. Tabarelli, C. Keitel, J. Gross, D. Baldauf

## Abstract

Successfully interpreting and navigating our natural visual environment requires us to track its dynamics constantly. Additionally, we focus our attention on behaviorally relevant stimuli to enhance their neural processing. Little is known, however, about how sustained attention affects the ongoing tracking of stimuli with rich natural temporal dynamics. Here, we used MRI-informed source reconstructions of magnetoencephalography (MEG) data to map to what extent various cortical areas track concurrent continuous quasi-rhythmic visual stimulation. Further, we tested how top-down visuo-spatial attention influences this tracking process. Our bilaterally presented quasi-rhythmic stimuli covered a dynamic range of 4 – 20Hz, subdivided into three distinct bands. As an experimental control, we also included strictly rhythmic stimulation (10 vs 12 Hz). Using a spectral measure of brain-stimulus coupling, we were able to track the neural processing of left vs. right stimuli independently, even while fluctuating within the same frequency range. The fidelity of neural tracking depended on the stimulation frequencies, decreasing for higher frequency bands. Both attended and non-attended stimuli were tracked beyond early visual cortices, in ventral and dorsal streams depending on the stimulus frequency. In general, tracking improved with the deployment of visuo-spatial attention to the stimulus location. Our results provide new insights into how human visual cortices process concurrent dynamic stimuli and provide a potential mechanism – namely increasing the temporal precision of tracking – for boosting the neural representation of attended input.

## 1. Introduction

Visual input from our environment conveys us with a massive stream of continuous information with rich temporal dynamics. Selective attention acts to filter this input in order to prevent distraction from our behavioral goals and optimally use the limited processing capacity of the brain. Attention filters out irrelevant information on various levels of processing and along various dimensions of sensory content, e.g., when selecting specific objects of interest (Duncan 1984, Baldauf & Desimone, 2014; Kim et al., 2017; Stoermer et al., 2019), when trying to extract relevant features (such as a specific color, Treue & Trujilio, 1999; Stoermer et al., 2014), or when focusing on particular spatial locations (*spatial attention*, see Mangun & Hillyard, 1991; Baldauf & Deubel, 2009). Spatial attention is of special importance for ubiquitous goal-directed interactions with our environment because it operates at the very core of efficient sensory-motor transformations, which we need to produce adaptive motor output. For example, in complex, real-world interactions, spatial attention regulates our ongoing eye movements (Moore & Armstrong, 2003; Baldauf & Deubel, 2008a), goal-directed reaches and grasps in the visual scene (Baldauf & Deubel, 2010; Baldauf et al., 2008b; Baldauf & Deubel, 2006), as well as generally navigating our environment while avoiding obstacles (Baldauf, 2018).

Our behavior therefore relies on constantly interpreting, updating and prioritizing the time-varying content, and thus the information embedded in the temporal structure of the continuous visual input stream. For instance, synchronicity of visual features contributes to the segmentation of scenes into objects and/or background (Alais, Blake, & Lee, 1998; Blake & Lee, 2000, 2005) and in extrapolating stimulus motion trajectories (Whitney, 2002) while compensating for neural latencies (Nijhawan, 1994). Consistency in the temporal structure of stimuli reduces reaction times and increases sensitivity to incoming stimuli by means of temporal expectation (Correa & Nobre, 2008). Schroeder and Lakatos (2009) suggested that this effect relies on a coupling of intrinsic brain rhythms to temporal regularities in sensory input and selective coupling might serve as a neural implementation of prioritizing (i.e. attending to) behaviorally relevant input. Conversely, our own locomotion may facilitate the processing of stationary scenes, as well as head or eye movements, where temporal dynamics of the resulting retinal representations can effectively contribute to high-acuity vision (Ahissar & Arieli, 2012; Schroeder, Wilson, Radman, Scharfman, & Lakatos, 2010). Taken together, these findings highlight the relevance of temporal dynamics in visual perception.

Despite the importance of temporal structure in the selective processing of dynamic visual input, neuroscientific research has historically focused on brain activity elicited as transient responses to briefly presented stimuli (Rust & Movshon, 2005). More recent studies have used continuous rhythmic stimulation, either to tag ongoing cortical visual processing (Andersen & Muller, 2010; Baldauf & Desimone, 2014), or to causally interfere with intrinsic brain rhythm involved in visual processing (Mathewson et al., 2012; Spaak, de Lange, & Jensen, 2014; Thut, Schyns, & Gross, 2011). These studies have used strictly rhythmic stimulation to facilitate the analysis of corresponding brain responses and/or maximise the impact on intrinsic brain activity. Thereby, they relied on stimulation in the form of periodic on-off flashes or sinusoidal modulations of the stimulus low-level features, such as colour or luminance. More recently, sinusoidal modulations were also applied to high-level, semantic content of visual objects by modulating their visibility through periodic phase-scrambling of image content without affecting low-level visual features (Baldauf & Desimone, 2014; Gordon, Tsuchiya, Koenig-Robert, & Hohwy, 2019). In all of these cases, strict rhythmicity has endowed stimuli with predictable and deterministic, yet also highly artificial temporal structure.

In our natural environment however, we mostly face more irregular and less predictable events and generalizing results from studies using strictly rhythmic stimulation to explain natural vision could be misleading (Blake & Lee, 2005; Haegens & Zion Golumbic, 2018). Recent research has therefore acknowledged the importance of temporal structure that carries only limited temporal regularity and can thus be considered quasi-rhythmic. In particular, brain activity couples to quasi-rhythmic natural stimulation during lip-reading and parsing hand gestures (Biau, Morís Fernández, Holle, Avila, & Soto-Faraco, 2016; Hauswald, Lithari, Collignon, Leonardelli, & Weisz, 2018; Park, Kayser, Thut, & Gross, 2016). The relevance of quasi-rhythmicity for visual perception has also been shown in monkey single cell recordings: Neurons in area MT and extrastriate cortex better discriminate concurrent quasi-rhythmic than constant-motion stimuli because they capitalise on the temporal fine structure of the visual input (Buračas, Zador, DeWeese, & Albright, 1998).

As for the neurophysiological basis of this tracking mechanism we note that visual cortex is abound with neuronal populations that detect features such as stimulus color, orientation, contrast and spatial frequency of incoming sensory information (Carandini, 2005). As shown by Buračas et al (1998), a given feature detector also represents the temporal dynamics of the stimuli through waxing and waning activity that corresponds to the intermittent occurrence of the preferred feature. The summed activity of all feature detectors over time can then be recorded as a macroscopic brain response with time-sensitive neuroimaging methods such as EEG/MEG. We consider this macroscopic signal reflecting the temporal dynamics of the stimulus as the *tracking* signal. A special type of tracking signal has been described before as consecutive transient responses to on/off rhythmic flicker stimulation (Capilla et al. 2010). Here we look into the more general case of tracking stimulus dynamics with only approximate rhythmicity and smooth continuous transitions in stimulus appearance. Also note that our notion of tracking is consistent with the recently proposed entrainment “in a broader sense”, i.e. measuring brain-stimulus synchronization in an oscillatory context but without assuming the explicit involvement of generators of intrinsic brain rhythms (Obleser & Kayser, 2019; for a similar point see Alexandrou, Saarinen, Kujala, & Salmelin, 2018).

To date, the cortical areas that respond to quasi-rhythmic visual stimulation have not been mapped in detail, nor has the attentional modulation of these processes been comprehensively described. Studies have identified a wide range of regions along the visual processing hierarchies that contribute to the generation of scalp level steady-state responses (SSRs) driven by strictly-rhythmic stimulation (Norcia et al., 2015; Parkkonen et al., 2008; Appelbaum et al., 2008; Di Russo et al., 2007; Fawcett, Barnes, Hillebrand, & Singh, 2004; Pastor, Artieda, Arbizu, Valencia, & Masdeu, 2003, Baldauf & Desimone 2014). These regions comprise early visual areas (V1-V4), as well as parts of the ventral (V8, fusiform & parahippocampal place areas) and dorsal streams (V3A) and motion sensitive areas (V5/MT). Cortical sources of SSRs can further differ between stimulus frequencies (M. A. Pastor, Artieda, Arbizu, Valencia, & Masdeu, 2003; M. Pastor, Valencia, Artieda, Alegre, & Masdeu, 2007). It follows that quasi-rhythmic, i.e. frequency-varying stimulation might drive a more complex pattern of cortical generators that similarly depends on the stimulation frequency band.

SSRs are also known to increase when the driving stimulus is attended indicating enhanced neural processing (reviewed in Vialatte, Maurice, Dauwels, & Cichocki, 2010; Norcia et al., 2015). This effect has been classically linked to a response gain mechanism (Müller, Teder-Sälejärvi, & Hillyard, 1998), and more recently to reflect greater synchronization between stimulus and brain dynamics (Gulbinaite, Roozendaal, & VanRullen, 2019; Joon Kim, Grabowecky, Paller, Muthu, & Suzuki, 2007; Keitel et al., 2019). Direct enquiries into which visual cortices show attentional modulation of SSRs have been limited. Electrical source imaging has implicated V1 consistently (Andersen & Muller, 2010; Keil et al., 2012; Keitel, Andersen, Quigley, & Müller, 2013, but see Hillyard et al., 1997) and other visual cortices, such as V4, the lateral occipital complex (LOC) and human area MT have been screened for attentional modulation after being pre-selected as regions of interest (Lauritzen, Ales, & Wade, 2010; Palomares, Ales, Wade, Cottereau, & Norcia, 2012). Studies investigating the cortical processing of naturally occurring quasi-rhythmic visual stimuli in low frequency bands (< 7 Hz), such as the tracking of a speaker’s lips movements (Hauswald, Lithari, Collignon, Leonardelli, & Weisz, 2018; Park, Kayser, Thut, & Gross, 2016) or hand gestures (Biau, Morís Fernández, Holle, Avila, & Soto-Faraco, 2016), have localized sources in circumscribed visual cortices without looking into detailed mapping of cortical regions and a modulation of the tracking response by visuo-spatial attention along the visual hierarchy.

Here, we aim to shed new light on how visual cortices track and prioritize rhythmic and quasi-rhythmic stimulation at task-relevant locations. We recorded MEG while participants viewed two stimuli, one positioned in the lower left and the other positioned in the lower right visual hemifield. Both exhibited well-defined contrast modulations at rates either varying within classical theta (4 – 7 Hz), alpha (8 – 13 Hz) and beta (14 – 20 Hz) bands, or with fixed frequencies of 10 Hz (left) and 12 Hz (right). Participants were cued on a trial-by-trial basis to focus on one of the two stimuli and to perform a target detection task at the attended position. An earlier investigation, recording EEG and using a similar paradigm, showed that brain responses follow the temporal evolution of the quasi-rhythmic stimulation precisely, thereby suggesting brain-stimulus coupling in all frequency bands (Keitel, Thut, & Gross, 2017). More specifically, spectral representations of the coupling indicated peaks within the stimulated frequency ranges. Theta- and alpha-band coupling further increased when the corresponding stimulus position was cued indicating that spatial attention biased brain-stimulus coupling.

The present investigation builds on these findings and extends them in several vital aspects. Most notably, we provide a detailed mapping of cortical areas that track visual stimulus dynamics based on MEG source reconstruction that use individual anatomical scans. We used a recently developed cortical parcellation that provides the most precise insights into the structural and functional organization of the human brain to date. This parcellation is based on a multi-modal atlas of the human brain, obtained by combining structural, diffusion, functional and resting state MRI data from 210 healthy young individuals (Glasser et al., 2016). We quantify the coupling of thus detailed cortices to the stimulation by means of spectral cross-coherence. First, we provide a proof of concept that our method can separate cortical responses to simultaneously presented stimuli oscillating within the same frequency band. Second, we investigate how temporal dynamics of quasi-rhythmic stimuli are tracked along the visual hierarchy and compare effects between frequency bands. Third, we investigate the effect of spatial attention, demonstrating that previously documented gain effects on SSRs (i.e. responses to strictly rhythmic stimulation) also exist for quasi-rhythmic stimuli, and exploit the fine-grained anatomical mapping to characterize these effects topographically for specific regions-of-interest in visual cortex.

## 2. Material and methods

### 2.1 Participants

24 healthy volunteers participated in the experiment. Participants were free of medication and with no history of neurological disease or injury. All of them reported right-handedness and normal or corrected-to-normal visual acuity and gave informed written consent prior to the recording session. After a preliminary inspection, we excluded data of seven participants (five because of the high number of eye movements during the experiment, one because of eye movements and contamination by severe myogenic artifacts and one because they were unable to perform the task as instructed). Finally, data from N=17 subjects (11 women, age 28.2 +/- 5 years, range 21-36 years) entered subsequent analyses. All procedures were approved by the University of Trento Ethics Committee.

### 2.2 Stimulation and task

Stimuli were generated using PsychToolbox (Brainard, 1997) and custom MATLAB code (The Mathworks, Natick, MA, USA) and projected on a translucent whiteboard using a ProPixx DLP projector (VPixx Technologies, Canada) at a 120 Hz refresh rate. The whiteboard, facing the participant frontally at a distance of 1 m (eyes-to-board), provided a physical projection area of 51×38 cm (width x height) and 1440×1080 pixel resolution. Stimulation (Fig. 1) consisted of two blurred checkerboard-like patterns (diameter 4° of visual angle) presented in the lower visual hemifields (horizontal/vertical distance from fixation 4° and 2.5° of visual angle). Two small concentric circles (maximum eccentricity 0.4°) were projected in the center of the screen and used as fixation and cue position. All stimuli were presented against a gray background (luminance 134 cm/m^2^). During each trial we modulated the Michelson contrast of the two patches continuously between a minimum of 10% (peak luminance 147 cd/m^2^) and a maximum of 90% (peak luminance 274 cd/m^2^), respectively. The underlying contrast modulation function was generated by frequency-modulating a carrier sinusoid with different modulation functions depending on the quasi-rhythmic stimulation condition (Fig 1), resulting in stimulation frequency bands of 4 - 7 Hz (*theta*, *θ*), 8 - 13 Hz (*alpha*, *α*) and 14 - 20 Hz (*beta*, *β*). In an additional condition, the steady state response condition (SSR), we used two strictly periodic contrast modulation functions of 10 and 12 Hz for the left/right patch, respectively. In the quasi-rhythmic conditions, each trial featured two different frequency modulation functions, one for each stimulus, generated by interpolating two different random uniform processes sampled at 500 ms intervals to the screen refresh rate of 120 Hz. Accordingly, the maximum velocity of frequency change was two full crossings of respective bandwidths per second. Moreover, we limited the Pearson’s correlation coefficient between the two frequency modulation functions to a maximum absolute value of 0.05 and generated a new set of functions for each trial. A trial started with a green hemi-circle presented for 500 ms between the two fixation circles and indicating the participant to attend the left or the right stimulus respectively (Fig 1). Following the cue, the two contrast modulated patches were presented for 3500 ms. Participants were instructed to detect occasional flashes that occurred at the cued position (*targets*) while ignoring flashes at the non-cued position (*distractors*). In order to generate the flashes, we replaced areas of the patch where the local contrast exceeded 10% of the background luminance with uniform grey at +/- 45% of the background luminance. Trials either contained two (1/6 of all trials), one (1/6) or no target/distractor events (2/3). Target/distractor events were randomly and equally distributed with respect to stimulus- and attended positions across the entire experiment. Each target/distractor was presented for 300 ms and the minimum interval between subsequent events within a trial was 800 ms. At the end of each trial the fixation circle turned red for 1 s, allowing subjects to blink. A new trial started after an inter-trial interval jittered between 600 ms and 1 s. The experiment formed a fully balanced design of the two independent manipulations stimulation condition (theta, alpha, beta and SSR) and attended position (left vs right) resulting in eight experimental conditions (e.g. theta-band stimulation – attend left). Practically, it consisted of eight blocks with 72 trials each, for a total of 144 trials per stimulation condition, randomly distributed across the eight blocks and with the cued position randomly drawn from an equal distribution within each condition’s set of trials. At the end of each block participants received on-screen feedback about average hit rate and reaction time. The average duration of a block was about 9 minutes.

**Fig. 1:**
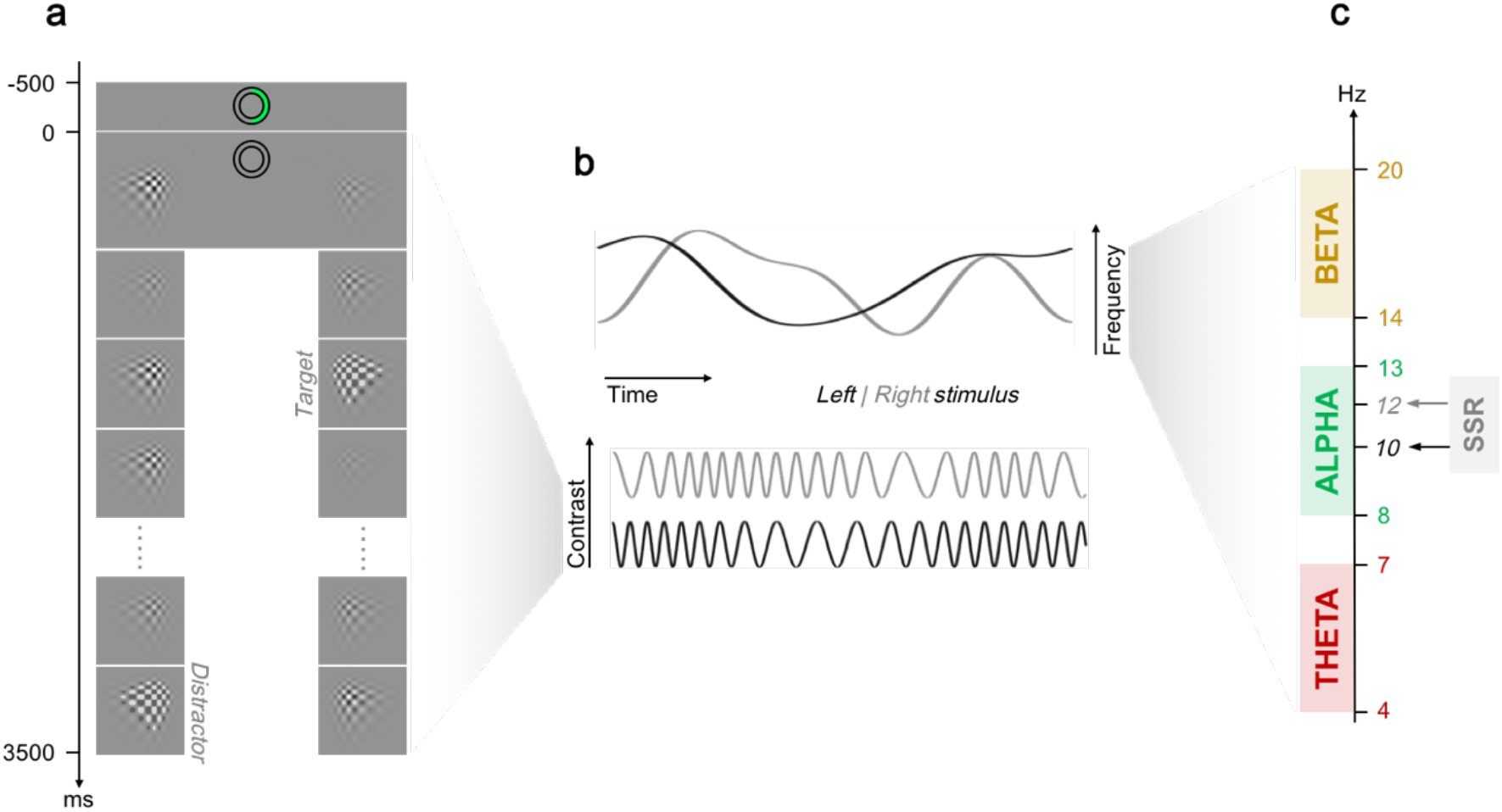
Details of the experiment and of the stimuli contrast modulation. (a) Trial time course: Central green left/right semi arc serves as attention cue for the first 500 ms. For the remainder of the trial the circle acts as fixation point while the two flickering stimuli are presented for 3500 ms. At random times, checkerboard contrast is flashed, producing a target or a distractor, depending on the initial cue. (b) Schematic illustration of stimulus dynamics: Stimulus contrast varies according to two uncorrelated frequency modulation functions within selected frequency bands with random trajectories in quasi-rhythmic stimulation conditions. (c) Depending on carrier frequency left/right stimulus contrast oscillates within theta, alpha or beta frequency band or with fixed frequencies of 10/12 Hz, respectively, in the SSR condition.

### 2.3 Procedure and data acquisition

All recording sessions took place during the day, between 9 am and 6 pm at the MEG Laboratory of the Centre for Mind/Brain Sciences of the University of Trento. After giving their informed consent, subjects were instructed about the task and the data acquisition. During instructions they performed a supervised training experimental block, in order to get acquainted with the task and as a final check that they correctly understood instructions. Before data acquisition we digitized the position of three anatomical landmarks (nasion and left/right auricle points) and five head position indicator (HPI) coils attached to the head, by using an electromagnetic position and orientation monitoring system (Polhemus Inc., Vermont, USA). Landmarks and HPI coils were digitized twice to reliably ensure a localization error of less than 1 mm. To improve the accuracy of co-registration with individual anatomies, we further digitized at least 330 additional points evenly spread out over the subject’s head and covering the nose profile. Participants performed the task seated comfortably in a two-layer magnetically shielded room (AK3B, Vacuum Schmelze, Hanau, Germany) where we recorded bio-magnetic activity from the brain using a Neuromag VectorView MEG scanner with 306 channels (204 first order planar gradiometers, 102 magnetometers, Elekta Inc., Helsinki, Finland). The MEG signal was sampled at 1 kHz, with a low pass antialiasing filter at 330 Hz and a high pass filter at 0.1 Hz. The position of the head during recordings was monitored at the beginning of each block. Responses were collected using a MEG compatible button response box (VPixx Technologies, Canada). During the experiment we recorded eye movements binocularly at 1 kHz with an eye-tracking system (SR Research Ltd., Ottawa, Canada). Moreover, at the onset of trial events (cue & stimulation onset, targets/distractors), a small white square was projected on the top left corner of the screen: the square was not visible for the participant but we used the corresponding signal, collected by a photodiode and a light-to-voltage converter (TKK Brain Research Unit), to correct for trigger delays introduced by the MEG build-in antialiasing filter, so as to reduce the overall temporal synchronization error to <1 ms.

### 2.4 Behavioral data analysis

Button presses less than 300 ms and more than 1100 ms after target/distractor onsets were excluded from the analysis, resulting in an average rate of discarded responses of 2.5 ± 0.6 % (median = 1.2 %). Due to their overall low occurrence we further excluded button presses recorded when no flash was presented at all, occurring at an average rate of = 5.8 ± 1.5 % (median = 2.3 %). We classified remaining valid responses, with respect to *target* and *distractors*, as defined by the cued position in each trial. In this context, hits and false alarms accounted for button presses in response to flash events that occurred at cued vs un-cued locations, respectively. We further defined correct rejections as omitted responses to a flash event at uncued location. Accuracy and false alarm rates are then defined as 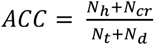 and 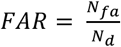 where *N_h_*, *N_cr_*, *N_fa_*, *N_t_* and N_d_ are the numbers of hits, correct rejections, false alarms, targets and distractors, respectively. For each participant we computed accuracy, false alarm rates and the median of reaction times in each of the eight experimental conditions. Accuracy and false alarms rates were subjected to a two-way repeated measures ANOVA with factors *stimulation condition* (theta, alpha, beta and SSR) and *cued position* (left vs right). The same analysis was performed on reaction times. To that end, each condition’s distribution of reaction times was centered with respect to the median reaction time across all conditions and condition-specific median RTs were derived from centered distributions for each participant. In all ANOVA analysis, when appropriate, we report the Greenhouse-Geisser corrected p-value (p_GG_) and epsilon (ε_GG_), together with original degrees of freedom and F-values. Effect sizes were evaluated by means of eta-squared η^2^ (Bakeman, 2005; Greenhouse & Geisser, 1959).

### 2.5 MEG data processing

MEG data were visually inspected and noisy channels were excluded from subsequent analyses. We filtered^©^ MEG data using the Neuromag MaxFilter implementation of temporally non-extended spatial Signal Source Separation (SSS) (Taulu, Simola, & Kajola, 2005; Taulu & Kajola, 2005), realigning each block sensor data to a reference frame. We chose the reference frame from all recorded head positions as the one with the minimal sum of Euclidean distances with respect to all other positions. On average, the movement compensated by SSS was 4 ± 2 mm and never exceeded 7.1 mm and the average residual rank of data after SSS was 69 ± 1 degrees of freedom. The center of the spherical harmonic expansion for SSS was selected by fitting a sphere on the points digitized on the participant’s head, excluding those below the nasion-auricular landmarks plane. A visual inspection of the data after SSS ensured the absence of signal-to-noise problems or sensor artifact residuals. Only data from planar gradiometers were included in further analyses. In a first step, data were scanned for artifacts. To this end, continuous data was filtered (1-99 Hz band pass filter; 49-51 Hz notch filter; Butterworth 4th order two-pass) and resampled to 200 Hz. After defining 4 s epochs starting 0.5 s before patch onset, we discarded all trials with blinks or loss of fixation (threshold at 2.5° of visual angle), according to eye-tracking data. Other epochs containing artifacts were manually marked for rejection. We further removed cardiac and other artifacts by inspecting independent components obtained from an extended Infomax ICA decomposition (Lee, Girolami, & Sejnowski, 1999). We set the maximum number of ICA components to be extracted as the residual degrees of freedom after SSS data rank reduction. This data inspection led to rejecting an average of 22 ± 14 % of trials and 3.6 ± 2 components for each subject, resulting in 110 ± 21, 110 ± 20, 113 ± 17 and 112 ± 20 average number of trials for the theta, alpha, beta and SSR conditions, respectively.

In a second pre-processing pass, continuous position-realigned and SSS-subjected data were bandpass-filtered between 3 – 21 Hz (Butterworth 4th order two-pass) encompassing the full bandwidth of visual stimulation across conditions, and then resampled to 500 Hz. After re-defining epochs as described above, previously marked trials with artifacts and components were removed from the filtered data. This was justified by the exchangeability of the order of ICA de-mixing and linear temporal filtering (Hyvärinen, Karhunen, & Oja, 2001) and by the spectral content of ICA removed artifacts being well below 99 Hz.

### 2.6 MEG source reconstruction based on individual anatomies

For each participant we collected a high-resolution T1-weighted anatomical scan using a 4 Tesla Bruker MedSpec Biospin MR scanner, with an 8-channel birdcage head coil (MP-RAGE; 1×1×1 mm; FOV, 256 x 224; 176 slices; TR = 2700 ms; TE = 4.18 ms; inversion time (TI), 1020ms; 7° flip angle). In order to consistently define coordinate systems, anatomical landmarks (anterior/posterior commissures and interhemispherical point) were marked on each subject’s brain scan. We obtained a mask for the volume enclosing the brain by segmenting T1 data with Fieldtrip (Oostenveld, Fries, Maris, & Schoffelen, 2011) and SPM software (Friston, 2007). The mask was visually inspected and then used in a standard FreeSurfer structural surface reconstruction pipeline (Dale, Fischl, & Sereno, 1999; Fischl, 2012; Fischl, Sereno, & Dale, 1999), so as to avoid potential issues in automatic skull-stripping. For each hemisphere, a high resolution (~160000 vertices) tessellated reconstruction of the external grey matter cortical surface was obtained, as well as a spherical inflated surface, whose original vertices grid was morphed to match anatomical landmarks on a template average of 40 subjects (Fischl, Sereno, Tootell, & Dale, 1999). The number of vertices on each cortical surface and co-registered sphere was consistently decimated to match the resolution of a 5th order recursively subdivided icosahedron by using MNE software (Hämäläinen, 2005) and custom MATLAB code. The resulting whole brain surface reconstruction (20484 vertices; 3.1 mm average source spacing) was used in modelling source positions for inverse solutions. In order to be able to perform group analysis at the source level, for each subject we computed an interpolation matrix between the decimated individual source model and the template average of 40 subjects. First the morphed participant’s co-registered spherical tessellation was aligned to the regularly spaced spherical inflation of the template cortical sheet. Then, for each point of the template sphere, we defined three linear interpolation coefficients as the inverse of the normalized distances from the vertices of the enclosing triangle in the subject coregistered sphere. Anatomical and MEG data were co-registered by manually matching digitized anatomical landmarks on the participant’s T1 scan. We further refined the co-registration by fitting additional digitized points to a 6000 vertices tessellation of the head surface, reconstructed from T1 data using Fieldtrip and SPM software. A realistic brain enclosing the tessellated surface (10242 vertices) was obtained from T1 data by mean of the watershed algorithm (Ségonne et al., 2004) and used to compute normalized lead fields with the FieldTrip single-shell method (Nolte, 2003). We carefully inspected and validated the result of each anatomical reconstruction and co-registration before further analysis. To reconstruct time series of neural activity at each source model position we used a Linear Constrained Minimum Variance (LCMV) approach (Van Veen, Van Drongelen, Yuchtman, & Suzuki, 1997). We computed a covariance matrix using data band pass filtered between 3 Hz and 21 Hz. The goal was to obtain spatial “common beamformers” allowing for the comparison of results between different flickering conditions, but still optimizing the solution with respect to the frequency bands of interest. Those common beamformers were then used to reconstruct dipole activity using data from each specific stimulation condition, further filtered in the correspondent contrast modulation frequency band (Butterworth 4th order two-pass, adding 1 Hz on both sides with respect to the stimulation frequency band). In computing the LCMV solution we applied a regularization of 1%, while choosing the optimal dipole orientation by mean of singular value decomposition. Finally, from time series of estimated neural activity, we redefined 1 s epochs starting from 0.5 s to 2.5 s after the patch onset with 50% overlap, resulting in 3 epochs per experimental trial. Furthermore, all segments containing target/distractors and/or responses were discarded. This led to an average number of epochs of 261 ± 50, 262 ± 48, 266 ± 38, and 264 ± 49 for the theta, alpha, beta and SSR condition, respectively. The ratio between attend left/right number of epochs was 1.08, 1.06, 1.06, and 1.03 for the theta, alpha, beta and SSR condition, respectively.

### 2.7 Individual coherence analysis

As a metric of coupling between neural activity at dipole *x* and the contrast modulation of the stimulus at position *s* we used spectral cross-coherence defined as:

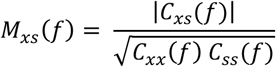

where *s* = *L* or *R* for *left* or *right* denote the corresponding stimuli. In the above formula, *C_ij_*(*f*) represents the cross spectral density at frequency *f*, averaged over epochs from Fourier coefficients *X* as follows:

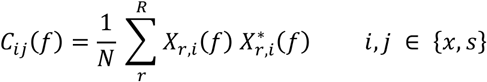

For strictly periodic stimulation, *C_ij_*(*f*) was estimated from Hanning-windowed Fourier coefficients and only the carrier frequency of the contrast modulation (*f* = 10 Hz and *f* = 12 Hz for left and right, respectively) was considered. Please note that, in this case, *M_xs_*(*f*) only depends on the phase consistency of the neural activity across trials and is thus equivalent to the classical inter trial phase coherence, as used in other studies on attentional modulation of visual steady state responses (Joon Kim, Grabowecky, Paller, Muthu, & Suzuki, 2007). For quasi-rhythmic conditions, we estimated *C_ij_*(*f*) using a Slepian multi-tapered approach (Percival & Walden, 1993), with epochs zero-padded so as to achieve a 0.5 Hz resolution. We chose the tapering bandwidth parameter so that the frequency smoothing interval spanned the whole frequency modulation band: in this way potential synchronization of the neural activity with the stimulus can be investigated by evaluating *M_xs_*(*f*) at the central frequency only (*f* = 5.5 Hz, *f* = 10.5 Hz and *f* = 17 Hz for the theta, alpha and beta conditions, respectively). Henceforth, for the sake of clarity, we will omit the frequency *f* in notation, as it is implicitly defined by the stimulation condition and stimulus position. In the quasi-rhythmic case we further computed a surrogate coherence 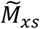 by using reversed contrast modulation functions. Time-reversing the stimulus had the benefit of keeping identical spectral properties while creating a signal that was not actually presented to the participant and in which the temporal information contained in the frequency modulation is destroyed. Evaluating *M_xs_* at each dipole *x* of the individual source model yielded a coherence map of the brain. We computed maps based on the cross-coherence with attended stimuli, with unattended stimuli and pooled over both attentional manipulations. Maps were evaluated for each stimulation condition separately. For each subject, we projected coherence maps from the individual source space {*x*}onto the template {*y*}using the correspondent pre-computed interpolation matrix, obtaining new maps *M_ys_* suitable for group analysis. Moreover, each map *M_ys_* was parcellated according to a multimodal brain atlas from the Human Connectome Project (Glasser et al., 2016). This mapping project combined structural, diffusion, functional and resting state MRI data from 210 healthy young adults to identify 180 regions of interest (ROIs), per hemisphere. For each of these ROIs *p* we extracted a single coherence value *M_ps_* as the 75^th^ percentile of the coherence distribution from dipoles *y* ∈ *p*. This procedure reduced sensitivity to outliers, while establishing a conservative threshold with respect to ROI overlap. Given the symmetry of the atlas, we further averaged coherence values over homologous ROIs contra/ipsilateral to the stimulus position, thus allowing us to represent contra/ipsi-lateral coherence as, by convention, right/left hemisphere in a set *M_p_* of 360 ROI values covering the whole brain.

### 2.8 Group analysis

Sample cross-coherence is a biased estimator. Therefore, before grand averaging and statistical testing, we stabilized the variance of the distribution of individual coherences using the Fisher transform (Enochson & Goodman, 1965). Grand averages over *N* subjects at each dipole/ROI were thus computed generically as

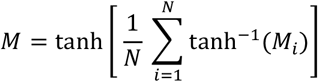

where the index *i* represents subjects. Moreover, before any group statistical analysis, we transformed single subject level coherences as *Z_i_* = tanh^−1^(*M_i_*).

Our first goal was to test whether cortical regions track the stimulus contrast modulation of both stimuli separately, also in the quasi-rhythmic conditions. The latter case is relevant as contrasts oscillate in the same frequency band and their contribution cannot be separated by conventional spectral decomposition (power spectra). We computed, for each flickering condition, dipole-wise grand averages of coherence with each stimulus using all trials, namely 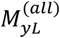 and 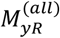 according to the above notation. We statistically compared them by running a two-tailed dependent-samples t-test under the null hypothesis that individual values 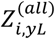 and 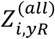 were drawn from the same distribution at each dipole. We used a non-parametric approach and a cluster-based correction (10000 permutations, alpha level = 0.05, cluster alpha = 0.05) to avoid false positives due to the high number of multiple comparisons (Maris, Schoffelen, & Fries, 2007).

As described above, we averaged coherence values of the same ROI from hemisphere contra/ipsi-lateral to the stimulus visual hemifield, thus obtaining parcellated maps *M_p_*. Before using these maps in subsequent analyses, we excluded any effect related to the stimulus position. To this aim, we statistically compared contra/ipsi-lateral coherence for left and right stimuli separately for each flickering condition and ROI *p*, testing whether 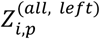 and 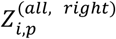 were drawn from the same distribution. We used a non-parametric two-tailed dependent samples t-test, where the distribution of the t-test statistics was evaluated with a permutation approach, using all permutations that can be drawn from a sample of 17 subjects (N=131072). The resulting 180 p-values – one per ROI – were Bonferroni-corrected for multiple comparisons because neighborhood between ROIs for a cluster-based correction cannot be meaningfully defined. The same conservative approach was used for all following ROI-based statistical analysis. Using this procedure did not reveal any significant effects of stimulus position therefore justifying a collapsing over 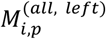 and 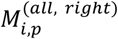 to yield a contra-/ipsilateral maps of cross-coherence.

The second goal of our work was to investigate how the temporal dynamics of quasi-rhythmic stimuli were tracked across cortical regions. For this reason, we compared ROI-specific coherence based on actually viewed stimuli 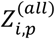 with surrogate coherence based on time-reversed stimuli 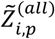. Under the null hypothesis of real and surrogate coherence belonging to the same distribution, we performed a one-tailed dependent sample t-test in a non-parametric permutation scheme, using all possible random permutations of labels *real* vs *surrogate* (N=131072); p-values resulting from the bootstrap procedure were Bonferroni corrected, considering 180 comparisons per hemisphere. Any region of interest rejecting the null hypothesis (Bonferroni corrected *p-value* < 0.01) was considered as tracking the stimulus dynamics. We repeated this analysis for each quasi-rhythmic flickering condition, obtaining three different sets of significant ROIs. In order to test whether coherence differences in significant ROIs between flickering conditions were due to sample variability, we performed a one-way repeated measures ANOVA with the factor *stimulation condition*(levels: theta, alpha, beta) on significant 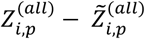 evaluating effect size as eta-squared *η*^2^ and applying a Greenhouse-Geisser correction for violation of sphericity, when appropriate (Bakeman, 2005; Greenhouse & Geisser, 1959). Resulting p-values were Bonferroni corrected (180 comparisons).

ROI reduction, and subsequent contra/ipsilateral collapsing might in principle mask small significant coherence effects, especially in regions that are not functionally lateralized. To rule out this possibility we ran an additional analysis, statistically comparing voxel-based coherence maps with surrogate data for each left/right cued condition separately, using for each stimulus those trials only in which it was attended (hereafter “*attended*” trials). We tested, in a permutation based one-tailed dependent sample t-test (N = 10000), whether 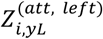 and 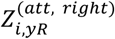 were drawn from the same distribution of 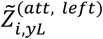 and 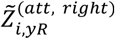, respectively. We performed a cluster-based correction for multiple comparison, using the sum of the cluster statistics in the permutation (Maris, Schoffelen, & Fries, 2007).

To validate our paradigm and analysis we further tested for known effects of attention on steady state responses (SSRs). Using our fine-grained spatial mapping approach allowed for an unprecedented level of detail which sub-regions of visual cortex were subject to an attentional bias and showed an increase in neural synchronization. To this purpose, we computed the difference of grand averages of parcellated coherence maps from *attended* and *unatten ded* trials 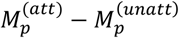, additionally testing if 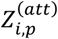 and 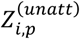 were different (for an alternative analysis using the Attentional Modulation Index – the AMI – see supplementary Fig S5). As above, we used a non-parametric permutation-based testing approach (all permutations) based on one-tailed dependent samples t-test and Bonferroni-correcting resulting 180 p-values. Using a one-tailed criterion was justified by a wealth of literature unequivocally showing a boosting effect of attention on coherence (Joon Kim, Grabowecky, Paller, Muthu, & Suzuki, 2007; Kashiwase, Matsumiya, Kuriki, & Shioiri, 2012; Porcu, Keitel, & Müller, 2013) and therefore allowing for a directed hypothesis.

We used the same approach to investigate attentional modulations in quasi-rhythmic flickering conditions. Here, we restricted the analysis of attentional gain effects in quasi-rhythmic conditions to the ROIs that were significantly different from surrogate coherence in at least one quasi-rhythmic flickering condition (Bonferroni corrected *p-value* < 0.01), i.e. those regions that were tracking the temporal dynamics of the stimuli. This led to evaluating attention effects in 56 and 21 areas for the contra- and ipsi-lateral hemisphere, respectively. Again, differences between the three quasi-rhythmic stimulation conditions might have been explained by sample variability. We tested this by performing a one-way repeated measures ANOVA on 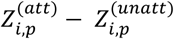 with factor *stimulation condition*, reporting eta-squared *η*^2^ and applying a Greenhouse-Geisser correction if required. Resulting p-values were Bonferroni corrected, considering 56 and 21 multiple comparisons for the contra- and ipsi-lateral hemisphere, respectively.

A comprehensive overview of all areas showing statistically significant effects is given in Tables 1 and 2. Name, description, and district (i.e. a superordinate grouping of neighboring cortices according to Glasser et al. 2016) are reported as well as MNI coordinates of the ROI centroid. Henceforth, for the sake of clarity, we will denote cross-coherence as XCOH and cross-coherence differences as ΔXCOH.

**Tab. 1:** List of ROIS, grouped by cortical district as from (Glasser et al., 2016); MNI coordinates of ROI centroid are reported in mm; within each group ROIs are ordered by increasing y-coordinate so that the first ROI in a district is the most posterior one.

| ROI | Description | District | ROI centroid in MNI coordinates (mm) |  |  |
| --- | --- | --- | --- | --- | --- |
|  |  |  | x | y | z |
| p24 | Area Posterior 24 | Anterior Cingulate Cortex | 3 | 35 | 16 |
| V3A | Area V3A | Dorsal Stream | 20 | -89 | 27 |
| V7 | Seventh Visual Area | Dorsal Stream | 26 | -80 | 35 |
| V6A | Area V6A | Dorsal Stream | 20 | -80 | 47 |
| V6 | Sixth Visual Area | Dorsal Stream | 18 | -76 | 32 |
| V3B | Area V3B | Dorsal Stream | 29 | -75 | 21 |
| IPS1 | Intra Parietal Sulcus Area 1 | Dorsal Stream | 23 | -68 | 39 |
| PGp | Area PGp | Inferior Parietal Cortex | 44 | -82 | 23 |
| IP0 | Area Intra Parietal 0 | Inferior Parietal Cortex | 33 | -73 | 29 |
| PGs | Area PGs | Inferior Parietal Cortex | 43 | -68 | 39 |
| IP1 | Area Intra Parietal 1 | Inferior Parietal Cortex | 33 | -66 | 43 |
| PGi | Area PGi | Inferior Parietal Cortex | 48 | -57 | 24 |
| PHT | Area PHT | Lateral Temporal Cortex | 59 | -55 | -4 |
| TE2p | Area TE2 Posterior | Lateral Temporal Cortex | 47 | -47 | -18 |
| TF | Area TF | Lateral Temporal Cortex | 41 | -23 | -27 |
| PHA3 | Para Hippocampal Area 3 | Medial Temporal Cortex | 35 | -40 | -15 |
| PHA1 | Para Hippocampal Area 1 | Medial Temporal Cortex | 23 | -38 | -16 |
| PHA2 | Para Hippocampal Area 2 | Medial Temporal Cortex | 33 | -35 | -16 |
| PreS | Pre Subiculum | Medial Temporal Cortex | 18 | -34 | -8 |
| V3CD | Area V3CD | MT+ Complex | 34 | -84 | 16 |
| LO2 | Lateral Occipital Area 2 | MT+ Complex | 46 | -84 | -6 |
| LO1 | Lateral Occipital Area 1 | MT+ Complex | 39 | -82 | 4 |
| LO3 | Lateral Occipital Area 3 | MT+ Complex | 48 | -77 | 12 |
| V4t | Area V4t | MT+ Complex | 47 | -77 | -1 |
| MT | Middle Temporal Area | MT+ Complex | 48 | -70 | 6 |
| MST | Medial Superior Temporal Area | MT+ complex | 45 | -66 | 3 |
| FST | Area FST | MT+ Complex | 46 | -64 | -4 |
| PH | Area PH | MT+ Complex | 47 | -63 | -11 |
| 5m | Area 5m | Paracentral Lobule | 4 | -42 | 63 |
| POS2 | Parieto Occipital Sulcus Area 2 | Posterior Cingulate Cortex | 9 | -71 | 36 |
| DVT | Dorsal Transitional Visual Area | Posterior Cingulate Cortex | 19 | -66 | 28 |
| 7m | Area 7m | Posterior Cingulate Cortex | 4 | -59 | 35 |
| POS1 | Parieto Occipital Sulcus Area 1 | Posterior Cingulate Cortex | 12 | -57 | 14 |
| v23ab | Area ventral 23 a+b | Posterior Cingulate Cortex | 3 | -52 | 20 |
| 31pd | Area 31pd | Posterior Cingulate Cortex | 12 | -52 | 38 |
| ProS | Pro Striate Area | Posterior Cingulate Cortex | 19 | -51 | 0 |
| 31pv | Area 31p Ventral | Posterior Cingulate Cortex | 9 | -47 | 33 |
| 31a | Area 31a | Posterior Cingulate Cortex | 6 | -44 | 37 |
| d23ab | Area Dorsal 23 a+b | Posterior Cingulate Cortex | 3 | -38 | 34 |
| RSC | Retro Splenial Complex | Posterior Cingulate Cortex | 4 | -36 | 27 |
| 23c | Area 23c | Posterior Cingulate Cortex | 12 | -31 | 41 |
| V3 | Third Visual Area | Primary and Early Visual Cortex | 29 | -91 | 8 |
| V4 | Fourth Visual Area | Primary and Early Visual Cortex | 33 | -85 | 1 |
| V1 | Primary Visual Cortex | Primary and Early Visual Cortex | 13 | -80 | 5 |
| V2 | Second Visual Area | Primary and Early Visual Cortex | 9 | -76 | -4 |
| 7PL | Lateral Area 7P | Superior Parietal Cortex | 13 | -71 | 56 |
| 7Pm | Medial Area 7P | Superior Parietal Cortex | 5 | -65 | 50 |
| MIP | Medial Intra Parietal Area | Superior Parietal Cortex | 22 | -65 | 46 |
| 7Am | Medial Area 7A | Superior Parietal Cortex | 8 | -58 | 61 |
| TPOJ3 | Area Temporo Parietal Occipital Junction 3 | Temporo Parieto Occipital Junction | 41 | -64 | 14 |
| TPOJ2 | Area Temporo Parietal Occipital Junction 2 | Temporo Parieto Occipital Junction | 52 | -59 | 8 |
| TPOJ1 | Area Temporo Parietal Occipital Junction 1 | Temporo Parieto Occipital Junction | 51 | -44 | 7 |
| PIT | Posterior Infero Temporal | Ventral Stream | 44 | -80 | -16 |
| V8 | Eighth Visual Area | Ventral Stream | 31 | -74 | -15 |
| VMV3 | Ventro Medial Visual Area 3 | Ventral Stream | 27 | -62 | -11 |
| FFC | Fusiform Face Complex | Ventral Stream | 39 | -57 | -22 |
| VMV1 | Ventro Medial Visual Area 1 | Ventral Stream | 20 | -55 | -12 |
| VMV2 | Ventro Medial Visual Area 2 | Ventral Stream | 29 | -53 | -8 |
| VVC | Ventral Visual Complex | Ventral Stream | 27 | -50 | -19 |

**Tab. 2:** List of ROI cortical districts, as from (Glasser et al., 2016), with abbreviations as used in the text; roman numerals in the last column are used in figures to identify the district.

| District name | District abbrev. | Roman numeral |
| --- | --- | --- |
| Primary and Early Visual Cortex | PEV | i |
| Dorsal Stream | DS | ii |
| Ventral Stream | VS | iii |
| MT+ Complex | MTC | iv |
| Inferior Parietal Cortex | IPC | v |
| Posterior Cingulate Cortex | PCC | vi |
| Lateral Temporal Cortex | LTC | vii |
| Medial Temporal Cortex | MTC | viii |
| Superior Parietal Cortex | SPC | ix |
| Temporo Parieto Occipital Junction | TPOJ | x |
| Other | OTH | xi |

### 2.9 MEG sensor based spectral and alpha lateralization analysis

In principle, participants could have solved the experimental task by using a global attention strategy: They could have attended to both stimuli simultaneously and, upon occurrence of a flash, decided ex post facto whether it fell on the cued or un-cued side. To control for this strategy, we conducted an additional analysis of alpha power spectral densities on the sensor level. The aim of this analysis was to reproduce the well described alpha lateralization effect linked to the deployment of spatial attention (Kelly, Lalor, Reilly, & Foxe, 2006; Thut, Nietzel, Brandt, & Pascual-Leone, 2006). In brief, attending to a spatial location in the right/left visual hemifield increases alpha power in ipsilateral visual cortices while alpha power in contralateral visual cortices decreases. Importantly, this alpha lateralization can be observed during states of sustained attention, i.e. while participants are expecting a stimulus to occur at the attended location.

To test for alpha lateralization, we Fourier transformed epochs starting from 0.25 s to 3.25 s after stimulus onset. For simplicity, this analysis was based on MEG gradiometer data only. Prior to spectral decomposition epochs were multi-tapered using Slepian sequences (Percival and Walden 1993) and then zero-padded to a length of 4 s, so as to achieve a spectral resolution of 0.25 Hz.

Using trials from all conditions, only separated by cued stimulus position (left vs right), we computed power spectral densities *P_x,s_*, where *x* denotes the sensor and *s* = *L* or *R* for (*left* or *right*) denote the corresponding cue. As a next step, we averaged *P_x,s_* over the alpha frequency band (8-14 Hz), for each participant and attend left/right conditions separately, obtaining averages 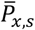. We then computed an alpha lateralization index from averaged alpha power spectral densities at each sensor position as 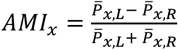 (Haegens, Handel, & Jensen, 2011; Thut, Nietzel, Brandt, & Pascual-Leone, 2006; Zumer, Scheeringa, Schoffelen, Norris, & Jensen, 2014).

Resulting AMI topographies of all participants were grand-averaged and tested against zero using two-tailed dependent sample t-tests in a non-parametric permutation approach as implemented in Fieldtrip (function ft_freqstatistics, using cluster correction for multiple comparisons with all possible permutations, N = 32768, alpha level = 0.05, cluster alpha = 0.05) and described by (Maris & Oostenveld, 2007).

Moreover, we computed sensor-based spectral profiles as well as cross-coherence with actual and surrogate (reversed) contrast modulation for all stimulation conditions, to allow for a comparison between an earlier EEG investigation (Keitel, Thut, & Gross, 2017) and the present data. In this analysis we pooled data from sensors in the back half of the helmet that pre-dominantly cover parieto-occipital brain areas including visual cortices.

Importantly, trials containing button presses and/or target/distractors were excluded from the alpha lateralization analysis. Thus, if participants followed the instructions, allocated their attention immediately after the cue and maintained this focus over the course of each trial we expected to see the typical alpha lateralization. Conversely, no alpha lateralization was to be observed in case of a global attention strategy.

## 3. Results

### 3.1 Behavioral results

Figure 2 shows results of behavioral analysis as outlined above, comparing performance measures across the four different stimulation conditions. Average accuracy was above 85% for all conditions (89 ± 3 % for *theta*; 88 ± 3 % for *alpha*; 85 ± 3 % for *beta*; 87 ± 3 % for *SSR, M ± SEM*) and the rates of false alarm were low (3.9 ± 1.5 % for *theta*; 2.7 ± 1.0 % for *alpha*; 1.7 ± 1.1 % for *beta*; 3.3 ± 1.4 % for *SSR*).

**Fig. 2:**
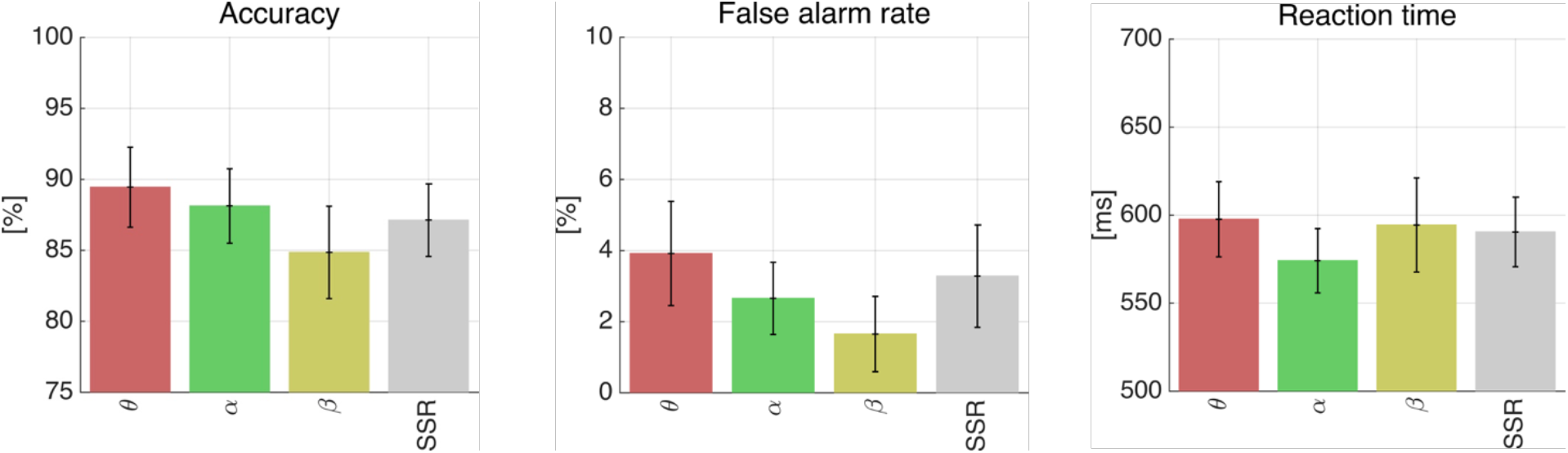
Behavioral results for all stimulation conditions. Error bars correspond to the standard error of the mean.

Testing for differences in performance between conditions, the stimulation frequency did not affect accuracy (F(3,48) = 2.4, p_GG_ = 0.12, ε_GG_ = 0.55, η^2^ = 0.02) and false alarm rates (F(3,48) = 1.2, p_GG_ = 0.33, ε_GG_ = 0.79, η^2^ = 0.03). Behavioral results further remained unaffected by the position of the attended stimulus (F(1,16) = 0.19, p = 0.67, η^2^ = 0.0005 for *accuracy*; F(1,16) = 0.86, p = 0.37, η^2^ = 0.004 and for *false alarm rate*), as well as the interaction of both factors, frequency range and stimulus position (F(3,48) = 0.37, p_GG_ = 0.74, ε_GG_ = 0.86, η^2^ = 0.003 for *accuracy;* F(3,48) = 1.1, p_GG_ = 0.36, ε_GG_ = 0.94, η^2^ = 0.013 for *false alarm rate;*).

Analyzing how fast participants responded to targets provided a similar pattern of results. Average reaction times were just below 600 ms in each condition (598 ± 21 ms for *theta;* 574 ± 18 ms for *alpha*; 594 ± 27 ms for *beta*; 590 ± 20 ms for *SSR*). Response speed did not change depending on the stimulation frequency (F(3,48) = 1.4, p_GG_ = 0.26, ε_GG_ = 0.59, η^2^ = 0.023, all tests based on centered reaction times), stimulus position (F(1,16) = 2.7, p = 0.12, η^2^ = 0.015) or a combination of both factors (F(3,48) = 0.4, p_GG_ = 0.71, ε_GG_ = 0.81, η^2^ = 0.015).

### 3.2 Sensor space: Brain-stimulus coupling and controlling for the sustained allocation of spatial attention

We used the well-known effect of alpha power hemispheric lateralization to test whether participants faithfully allocated their attention to the stimulus location as indicated by the cue in the beginning of each trial. Therefore, we conducted spectral analyses in sensor space. We found that brain activity picked up with MEG gradiometers was dominated by a prominent alpha band response (~10 Hz peak) during stimulation, irrespective of the stimulation frequency (Figure 3a). Sensor space analysis further demonstrated stimulus-specific coupling of brain responses as measured by spectral cross-coherence for strictly rhythmic and quasi-rhythmic stimulation (XCOH, Figure 3b). Note that reversing contrast modulation functions to conduct a surrogate analysis did not reveal any cross-coherence (XCOH) between brain response and stimulus. Finally, analysis of average alpha power (Fig 3c) showed two lateralized clusters in which the attentional modulation index (AMI) differed from zero (left: T_sum_ = 34.4, p = 0.015; right: T_sum_ = −19.7, p = 0.03). This result confirmed that participants allocated sustained visuo-spatial attention to the cued position throughout the stimulation period and thus were following instructions as intended. Conversely, this finding rules out an alternative global attention strategy that participants could have adopted to solve the task.

**Fig. 3:**
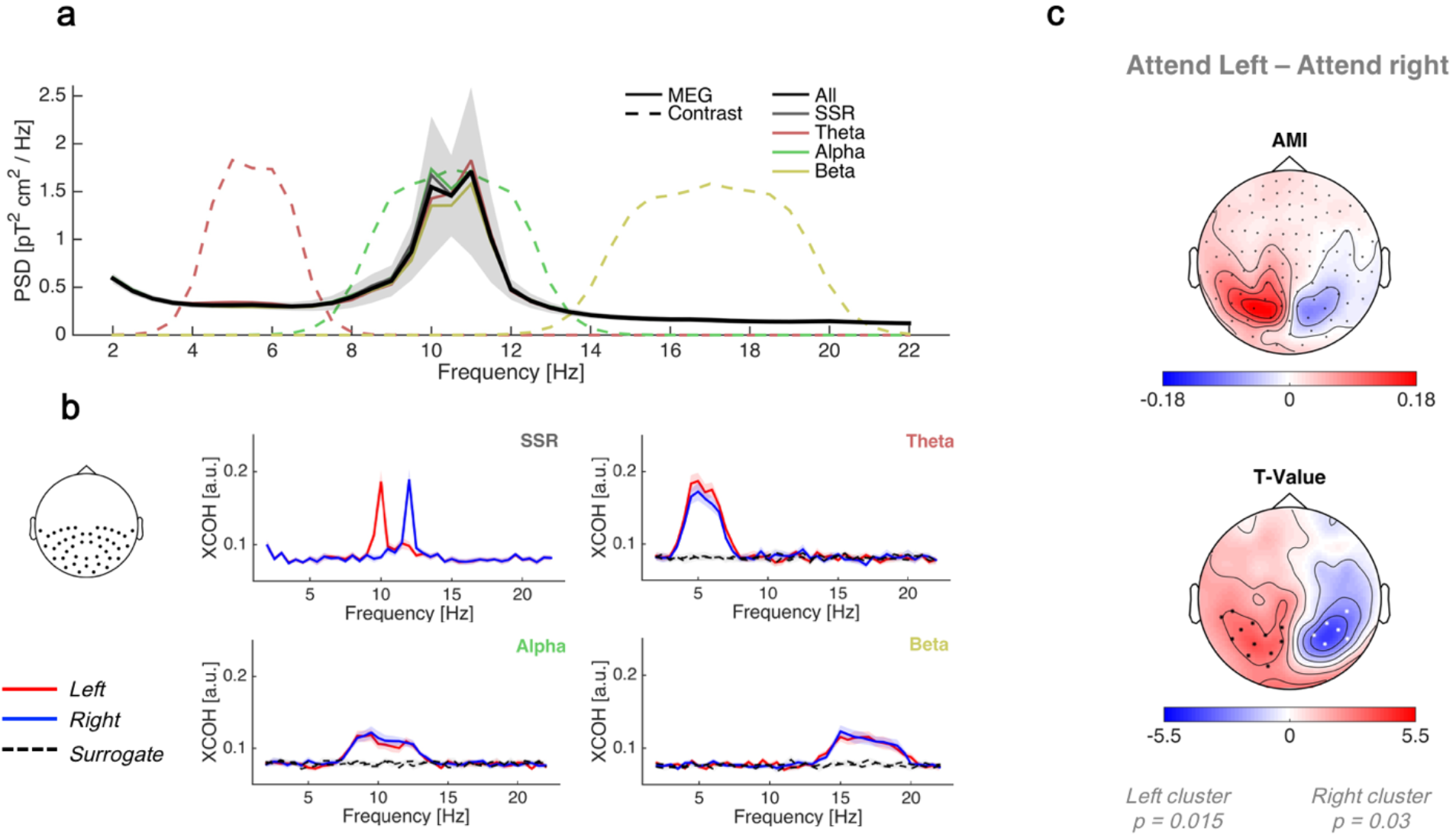
Sensor based spectral analysis. a) Power spectral densities of MEG sensors, averaged over all subjects for the four stimulation conditions separately (colored lines) and pooled across conditions (black line). Dashed lines represent scaled spectral representations of the stimulus contrast modulations in the quasi-rhythmic conditions. b) Cross–coherence (XCOH) between sensors for real (red and blue) and surrogate (reversed) contrast modulations (dashed lines) averaged across MEG gradiometers in the posterior hemisphere of the helmet (see topography). c) Scalp maps of the attention modulation index (AMI) of alpha power and correspondent statistical analysis show the typical lateralization effect during sustained focused spatial attention.

### 3.3 Spatial separation of cortical stimulus processing

Grand-average cortical maps of neural coherence with left and right stimuli (Fig. 4) demonstrate that our approach can reliably separate stimulus-specific neural activity driven by two simultaneously presented dynamic stimuli for SSR and quasi-rhythmic conditions alike. More specifically, we found phase synchronization (XCOH) maxima with each stimulus in the occipital areas within the respective contralateral visual cortex (Fig 4a). XCOH was of comparable magnitude in SSR and *theta* conditions, while decreasing in *alpha* and further in *beta*. This was confirmed by a lateralization analysis based on dipole-wise statistical tests of the difference in the spatial distribution of coherence with the left minus the right stimulus (Fig. 4b). Although neural tracking was stimulus-specific within the calcarine sulcus (see medial views to the left and right in Figure 4b) the center panels (occipital view) of statistical parametric brain maps show a reduced specificity on the gyri immediately surrounding the calcarine sulcus outside the longitudinal fissure. Starting from these extrastriate early visual cortices, tracking specificity increased towards downstream visual areas. Also, coherence maps in Fig. 4a show a spread from the contralateral to the ipsilateral hemisphere. This is most likely due to the anatomical separation being close to the limit of beamforming resolution. Another possibility are errors in segmenting ventral occipital cortex by using T1-weighted MR anatomical scans that might introduce spread effects (Winawer, Horiguchi, Sayres, Amano, & Wandell, 2010). No tracking effect was found for frontal regions when controlling for lateralization. For quasi-rhythmic stimulation conditions, this result was confirmed in a control analysis in which the cross-coherence with actual and surrogate (reversed) contrast modulation was statistically compared at the dipole level without collapsing into ROIs, for left and right hemispheres separately and using those trials only in which the respective stimulus was attended. This analysis approach particularly aimed at exploiting possible small coherence effects in frontal areas that do not necessarily underlie a spatiotopic lateralization. Results of this analysis largely corroborated our primary finding, mostly showing stimulus tracking within occipital areas (Supplementary Fig 4), with the notable exception of a cluster in the temporal lobe when tracking the left stimulus in the alpha condition. The specificity and location of this effect makes an interpretation highly speculative. Moreover, as mentioned in the Methods section, the statistical analysis level excluded effects of stimulus position on per-ROI XCOH. We took these results as a justification to collapse per-ROI XCOH across left and right stimuli in subsequent analyses, while converting data to a contra-vs ipsilateral representation of respective stimulus tracking.

**Fig. 4:**
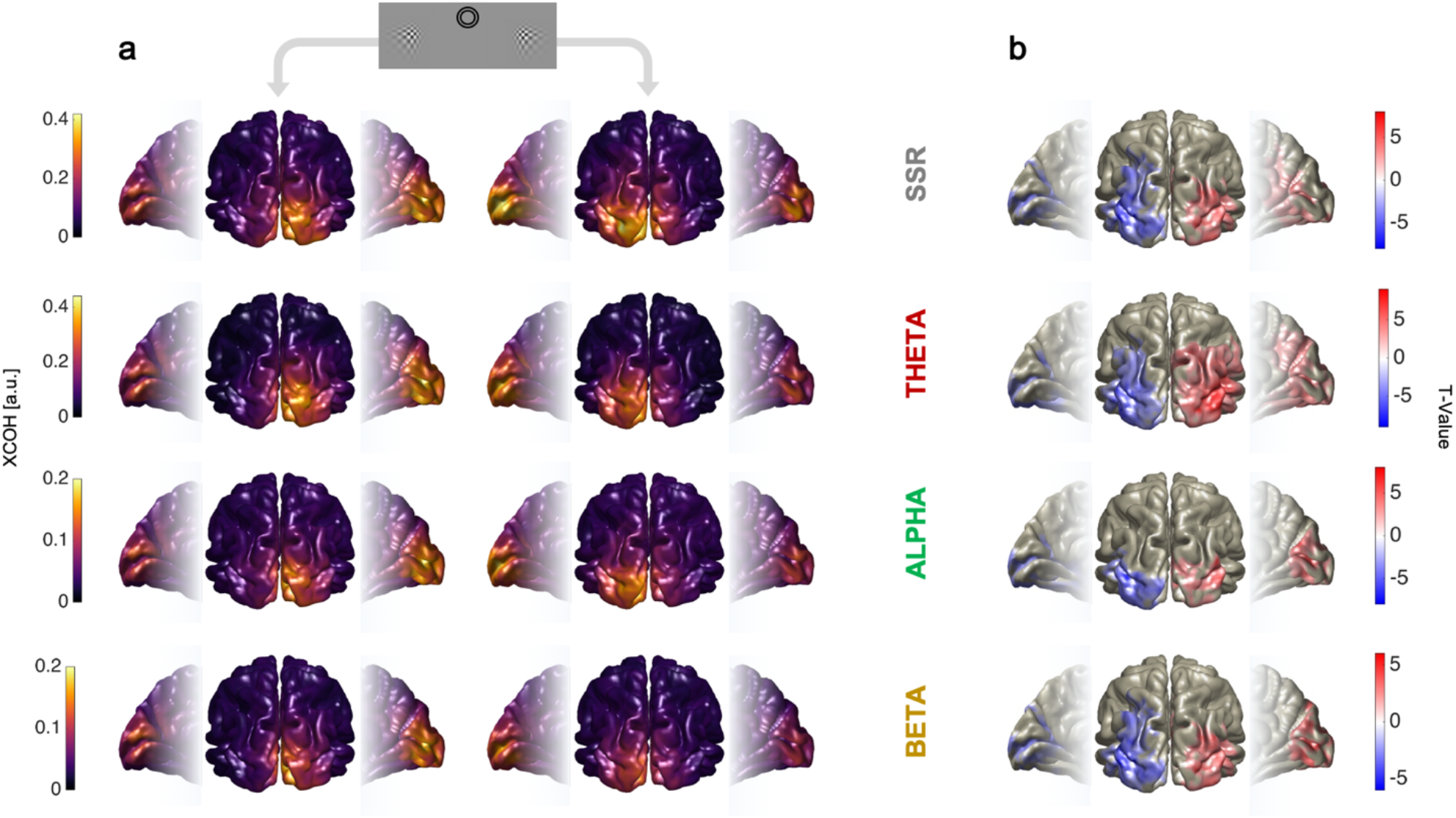
Spatial separation of cortical stimulus processing. (a) Dipole-wise cross-coherence (XCOH) with left 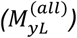 and right 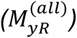 stimulus contrast in all stimulation conditions, collapsed across trials of attended and unattended conditions, respectively. In (b): Maps of t-values obtained by a two-sided non-parametric dependent samples t-test comparing Fisher stabilized XCOH with left 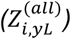 and right 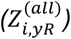 stimulus contrast (alpha level = 0.05, cluster alpha = 0.05).

### 3.4 Cortical tracking of underlying temporal dynamics

Mostly contralateral occipital visual areas successfully tracked temporal information contained in the quasi-rhythmic stimulus contrast modulation (Fig. 5). Intensity of the tracking hereby crucially depended on the stimulation frequency. During slow theta modulations XCOH propagated the furthest along the visual processing hierarchy and produced greatest XCOH. Specifically, primary and early visual areas (V1, V2, V3 and V4), as well as some posterior areas of the dorsal stream (V3A and V6), of the ventral stream (V8 and PIT) and of the MT+ complex (V3CD, LO1, LO2, LO3 and V4t) consistently showed phase coherence with the temporal dynamics of the stimuli for all quasi-rhythmic conditions. The extent to which quasi-rhythmic frequency modulations were tracked depended on the stimulation frequency band in all areas but TE2p, PFT, TF, PHA1 and 5m, as demonstrated by the results of ROI-specific ANOVAs (black asterisks in the boxplots in Fig. 5b; also see map of effect sizes in Fig.5c).

**Fig. 5:**
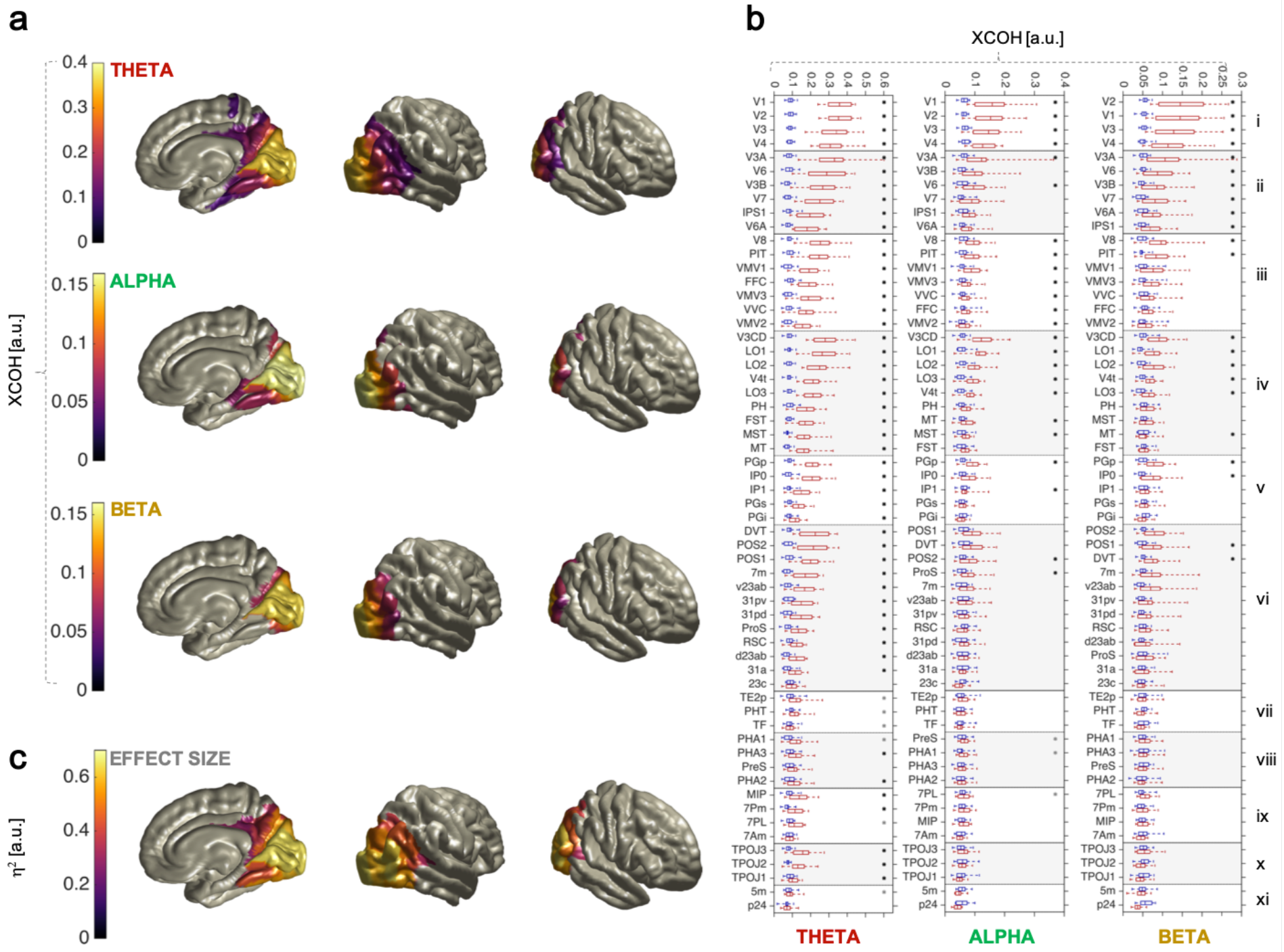
Contralateral cortical tracking of underlying quasi-rhythmic temporal dynamics. In (a) grand-average ROI-specific cross-coherence (XCOH) with contralateral stimulus 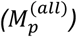 is reported for all quasi-rhythmic stimulation conditions; each cortical ROI is colored according to its correspondent value; data collapsed across attention conditions; ROIs are masked by significance (Bonferroni corrected p-value < 0.01, 180 comparisons) from a one-sided non-parametric dependent samples t-test comparing Fisher stabilized XCOH with actual 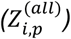 and surrogate 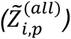 contrast modulations. Asterisks on boxplots in (b) indicates areas significantly tracking the contrast underlying dynamics according to the above mentioned statistical comparison; boxplots show XCOH with actual (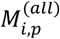 in red) and surrogate contrast (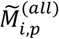 in blue) for each ROI in Table 1; roman numerals identify ROI districts, as per Table 2; black asterisks indicate that the magnitude of XCOH further depends on the stimulation frequency, as determined by a one-way repeated-measures ANOVA on Fisher stabilized XCOH differences 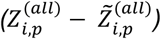 between actual and surrogate data. Corresponding effect sizes are reported in (c) only including ROIs whose XCOH depended on the stimulation frequency. Stimulus frequency has largest effects on XCOH in early visual cortices.

Except for posterior occipital areas outlined above, there were differences in how the temporal dynamics information was propagated along the visual streams. Boxplots in Fig.5b) show that the dorsal stream was consistently involved in tracking *theta* and *beta*, but not *alpha* dynamics. A different pattern emerged for the ventral stream, which was more sensitive to *theta* and *alpha* temporal dynamics, while being almost insensitive to *beta*. Results for the MT+ complex were less clear-cut, despite a systematic stimulus-frequency dependent decrease in the number of sensitive ROIs was observed. For the *theta* condition, temporal dynamics information propagated beyond classical visual districts, into the Posterior Cingulate Cortex (PCC), Inferior and Superior Parietal Cortex (IPC and SPC), Medial Temporal Cortex (MTC) as well as to the Temporo Parietal Occipital Junction. ANOVAs confirmed that ROI sensitivity in these districts depend on the stimulation frequency and is mostly absent in *alpha* and *beta* conditions, except for some posterior occipital areas. Further areas, e.g. within the Lateral Temporal Cortex (LTC), showed significant coherence with theta stimulation. A cortical representation of effect sizes (η^2^), derived from ROI-specific ANOVAs (Fig 5c), illustrates how much the magnitude of XCOH depended on the factor of stimulus frequency (*theta*, *alpha*, *beta*). Ipsilateral areas (Supplementary Figure 1) generally showed weaker tracking of temporal dynamics and were largely restricted to those regions that extend into the longitudinal fissure. Notably, a selection of ipsilateral areas outside the longitudinal fissure also tracked the temporal dynamics (V3A, V4, V8, IPS1, PIT). Again, XCOH magnitude depended on the stimulation frequency in some cases. It is worth noting that neither analysis implicated frontal areas in the tracking of quasi-rhythmic stimulation. This last result was confirmed in an additional control analysis in which the XCOH with actual and surrogate (reversed) contrast modulation was statistically compared at the dipole level without collapsing into ROIs, for left and right hemispheres separately and using those trials only in which the respective stimulus was attended (Supplementary Fig 4).

### 3.5 Attentional modulation of Steady State Responses

Our results confirmed previous findings that attention modulates the steady state response within visual cortices contralateral to the driving stimulus (Fig. 6). Primary and early visual cortices showed high attentional modulation (ΔXCOH = 0.075 for V1 and ΔXCOH > 0.063 for V2, V3 and V4) as well as their anterior neighbors POS1 (ΔXCOH = 0.061), V6 (ΔXCOH = 0.062). Note that V8 showed numerically greater attentional modulation (ΔXCOH =0.078), surpassing the attentional modulation in V1 and in its neighbor V4. Despite being anatomically distant from occipital visual cortices, areas 7Am in the Superior Parietal Cortex and p24 in the Anterior Cingulate Cortex also showed attentional modulation for the steady state response (ΔXCOH = 0.051 for 7Am and (ΔXCOH = 0.047 for p24).

**Fig. 6:**
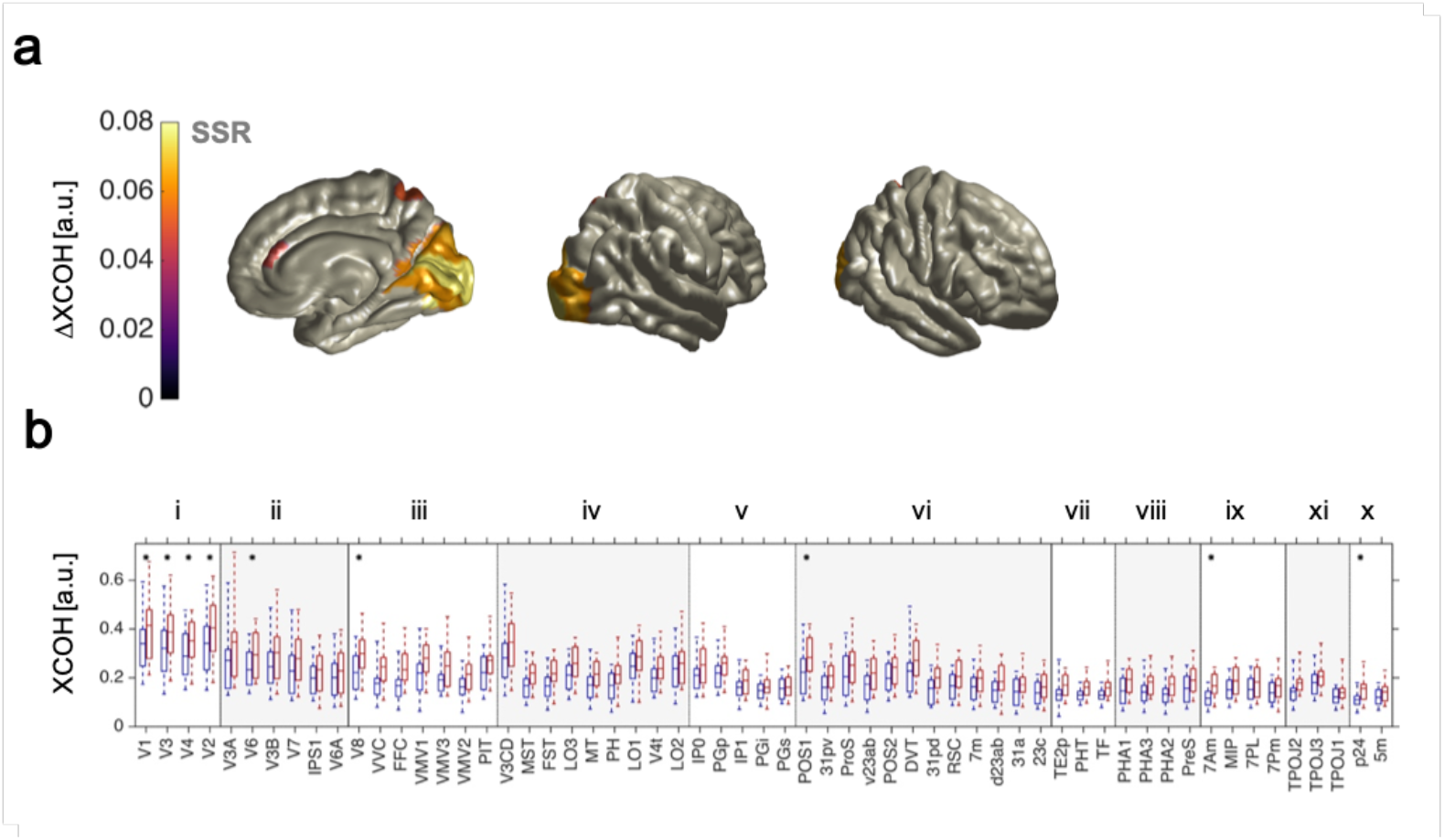
Contralateral attentional modulation of Steady State Responses. In (a) the difference 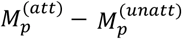 of grand-average ROI-specific cross-coherence (XCOH) with a contralateral attended and unattended stimulus is shown; each ROI on the cortex is colored according to the difference in XCOH (= ΔXCOH); only significant differences are displayed (Bonferroni corrected p-value < 0.05, 180 comparisons), according to a one-sided non-parametric dependent-samples t-test comparing Fisher stabilized XCOH in the attended 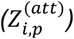 and unattended 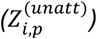 condition. Asterisks on boxplots in (b) indicates areas where the XCOH with the contrast modulation is significantly enhanced as per the above statistical comparison; boxplots show XCOH in the attended (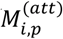 in red) and unattended (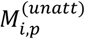 in blue) condition for each ROI in Table 1; roman numerals identify ROI districts, as from Table 2.

Regarding attentional modulation in ipsilateral cortices we found weaker and less widespread effects (Supp. Fig. 2). These effects were only observed for areas that were directly adjacent homologues of contralateral areas within the longitudinal fissure that also showed attention effects (ΔXCOH = 0.061 for ipsilateral V1; ΔXCOH = 0.048 for ipsilateral V2; ΔXCOH = 0.043 for ipsilateral p24). It is thus possible that these effects reflect the spatial blurring of the beamformer rather than a genuine effect, or ventral T1 segmentation errors as reported in (Winawer, Horiguchi, Sayres, Amano, & Wandell, 2010).

### 3.6 Attentional modulation in quasi-rhythmic conditions

Similar to the SSRs, attention enhanced cross coherence between stimulus contrast and brain responses in the quasi-rhythmic conditions (Fig. 7). Primary visual area V1 showed attentional tracking enhancement consistently across all quasi-rhythmic stimulation conditions. The magnitude of this effect did not depend on the stimulation frequency (F(2,32) = 8.1, p_GG_ = 0.39 Bonferroni corrected for 56 comparisons, ε_GG_ = 0.63, η^2^ = 0.24). The attentional modulation effect was spatially most confined in the *alpha* condition, where V1 was the only region to show significant modulation (ΔXCOH = 0.027). For *beta*-band stimuli, V2 (ΔXCOH = 0.017), V8 (ΔXCOH = 0.025) and V3A (ΔXCOH = 0.028) showed additional modulations whereby gain effects in V3A and V8 numerically surpassed V1 (ΔXCOH = 0.018) and V2 effects.

**Fig. 7:**
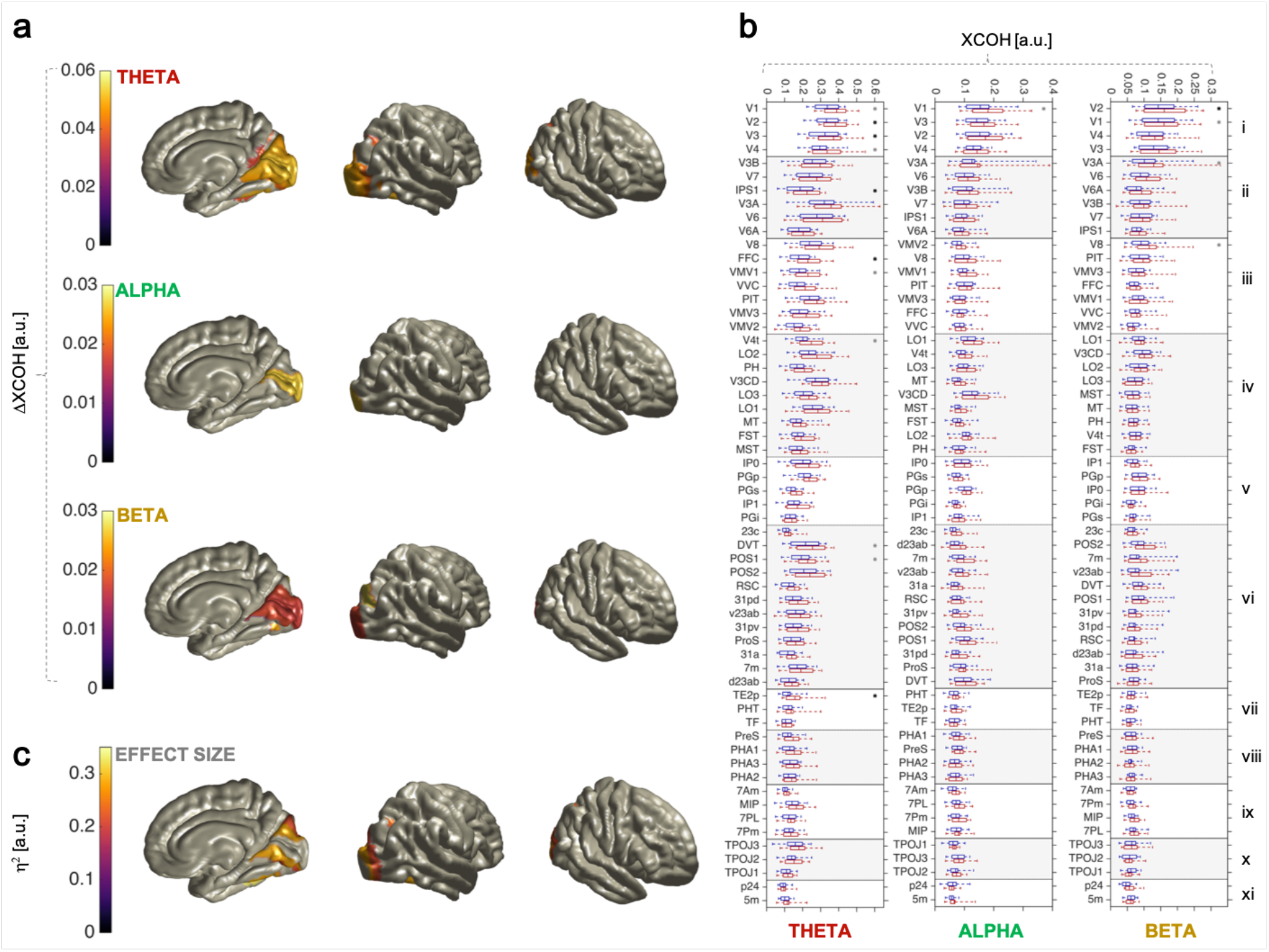
Contralateral attentional modulation in quasi-rhythmic conditions. In (a) the difference 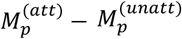 of grand-average ROI-specific cross-coherence (XCOH) with contralateral attended and unattended stimulus is shown; each ROI on the cortex is colored according to the difference in XCOH (= ΔXCOH); only significant differences are displayed (Bonferroni corrected p-value < 0.05, 56 comparisons), according to a one-sided non-parametric dependent-samples t-test comparing Fisher stabilized XCOH in the attended 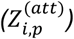 and unattended 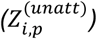 condition. Asterisks on boxplots in (b) indicates areas where the XCOH with contrast modulation is significantly enhanced as per the above statistical comparison; boxplots show XCOH in the attended (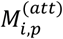 in red) and unattended (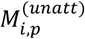 in blue) condition for each ROI in Table 1; roman numerals identify ROI districts, as per Table 2. Black asterisks indicate that the difference also depends on the stimulation frequency, according to a one-way repeated-measures ANOVA on Fisher stabilized ΔXCOH 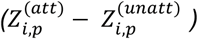. Corresponding effect sizes are reported in (b) only including ROIs whose attention effect (ΔXCOH) depended on the stimulation frequency. Stimulus frequency has largest effects on ΔXCOH in early visual cortices excluding V1.

In the *theta* condition we observed the overall biggest and spatially most widely distributed attentional modulation. Attention modulated XCOH mostly in primary and early visual areas (ΔXCOH = 0.052 for V1; ΔXCOH = 0.05 for V2; ΔXCOH = 0.048 for V3; ΔXCOH = 0.041 for V4), as well as neighboring areas in the PCC (ΔXCOH = 0.036 for POS1) and in the Ventral Stream (ΔXCOH = 0.048 for VMV1). For the ventral stream, we found attentional modulation in the Fusiform cortex (FFC) and area V4t despite their anatomical distance from early visual cortices. Notably, the FFC gain effect (ΔXCOH = 0.051) was numerically comparable to the V1 effect and the effect in area V4t was of similar magnitude as those in early visual areas V2-V4. Following the ventral stream, area TE2p, an area adjacent to FFC, showed further attentional modulation (ΔXCOH = 0.037). Dorsal stream areas exhibited attentional gain effects as well in IPS1 (ΔXCOH = 0.039) and in its ventral neighbor, the Dorsal Transitional Visual area DVT (ΔXCOH = 0.041).

During theta stimulation we also observed attentional modulation of XCOH in areas of the ipsilateral hemisphere (Supp. Fig. 3). This was mostly limited to primary and early visual areas (ΔXCOH = 0.037 for V1; ΔXCOH = 0.036 for V2; ΔXCOH = 0.024 for V3) and in proximal area ProS within the longitudinal fissure (ΔXCOH = 0.03). Similar to the contralateral hemisphere, gain effects were also found in the ventral stream in adjacent areas V8 (ΔXCOH = 0.032) and PIT (ΔXCOH = 0.032).

## 4. Discussion

In the present study, we source-localized MEG responses to bilaterally presented, competing quasi-rhythmic visual stimuli. Using a novel detailed anatomical parcellation of the cortex (Glasser et al. 2016) we identified regions that tracked the temporal evolution of both stimuli. Additionally, we measured effects of spatial attention on the tracking process by cueing participants to perform a detection task on the left or right. Large parts of visual cortex showed systematic coupling with the stimulus. This coupling was stronger in contralateral cortices (within the limits of the spatial uncertainty of the inverse model solution), consistent with the spatiotopic projection of sensory input into early visual cortices. This lateralization further indicated that coupling occurred in a stimulus-specific fashion although contrast fluctuations occurred within the same frequency range for stimuli on both sides. Among all quasi-rhythmic stimulation conditions, *theta*-band stimulation produced the strongest and cortically most widespread coupling. Attention to the stimulus location increased the synchronization with both strictly- and quasi-rhythmic stimuli in primary and early visual cortices and beyond. Cortical patterns of attentional modulation differed between stimulation conditions with *theta* showing the most extensive spread of gain effects.

### 4.1 Tracking of quasi-rhythmic temporal dynamics along the visual processing hierarchy

Our results corroborate earlier results of an EEG investigation (Keitel, Thut, & Gross, 2017). Using a similar paradigm, the authors reported that scalp-level EEG indicated phase-locking to quasi-rhythmic stimulation with a similar gradient from strong, widespread *theta* tracking to relatively weak and more confined *beta* tracking. Here, we extend these findings with a spatially detailed characterization of the phenomenon on the cortex. We present a detailed and comprehensive cortical map of how the brain tracks quasi-rhythmic temporal dynamics of visual input. Crucially, this map elucidates how different parts of visual cortex contribute to the processing of continuous dynamic stimuli that possess a higher degree of ecological validity than static transient or strictly rhythmic stimulation (Blake & Lee, 2005; Haegens & Zion Golumbic, 2018).

We found that a range of occipital, parietal and temporal brain regions tracked the temporal dynamics underlying contrast modulations during *theta* (4 – 7Hz), *alpha* (8 – 13 Hz) and *beta*-band (14 – 20 Hz) stimulation. Predominantly regions in the cerebral hemisphere *contralateral* to the spatial position of the stimulus produced responses that were strongly phase-locked to the contrast modulation of the stimuli irrespective of the stimulation frequency we observed primary and early visual areas (V1 – V4), as well as adjacent areas (mainly ventral and dorsal streams, MT+ complex).

Stressing the role of motion-processing areas in tracking the temporal structure of time-varying stimuli, our findings involved the MT+ complex in all stimulation conditions, albeit with decreasing strength and extent with increasing frequency band. The general involvement of the MT+ complex is consistent with findings by Buračas et al. (1998), who measured Local Field Potentials in area MT, the monkey homologue of human MT+ cortex (Orban, Van Essen, & Vanduffel, 2004), to demonstrate the superiority of quasi-over strictly rhythmic stimulation in a discrimination task. As in the present study, they implemented quasi-rhythmicity as fluctuations in stimulus timing in the range of 30-300 ms, roughly corresponding to the full range covered by our stimulation conditions (4 – 20 Hz).

We also observed differences in the extent of where and how far cortical tracking propagated in the cortex: *theta* tracking was strongest overall and exceeded *alpha* and *beta* tracking by propagating into lateral and medial temporal cortices, superior parietal cortex and the temporo-parietal junction. This prominence of low-frequency stimuli in terms of neural tracking could have several reasons: assuming some constant temporal variation (jitter) at each synaptic transmission to the subsequent cortical area, the neural synchronization with slower-paced stimulation in higher-level areas would suffer less compared to fast stimulations. A second reason could be the pre-dominance of a specific frequency band, like theta, as an evolutionarily established communication channel. This latter reason for the relatively strong propagation of theta-band stimulations seems in accordance with findings of research into the neural underpinnings of speech-reading and hand gestures. The frequency range of lip movements and gestures stimuli (1-8 Hz) is broadly comparable with our *theta* stimulation. Park et al (2016) reported tracking of lip movements in primary and early visual areas up to occipito-parietal areas when lip movements were incongruent with (i.e. unrelated to) concurrently presented speech (this situation resembled our experimental setup most closely because we did not present auditory stimuli). Also, Hauswald et al. (2018) reported that ventral and dorsal occipital areas, including primary and early visual cortices, follow the dynamics of lip movements in the absence of audible speech. Using Granger causality to test functional connectivity, they further found evidence for top-down control over early visual areas, suggesting the involvement of a brain-wide network in processing visual lips movements with the aim of speech intelligibility. Biau et al (2016) used a spatially more detailed fMRI approach to study gesture processing. They found that quasi-rhythmic hand gestures elicited broad occipital activation. This activation extended to the Medial Temporal Gyrus and the Superior Temporal sulcus, partially overlapping with our results for *theta* tracking.

In contrast to the tracking of slow (theta) dynamics the cortical sources of the tracking of quasi-rhythmic alpha and beta band stimulation have not been described before to our knowledge. The present results indicate that dorsal and ventral streams track the temporal dynamics differently in these two conditions, whereas both streams are involved in tracking *theta* range dynamics. Specifically, ventral stream areas are more involved in tracking dynamics in the 8-13 Hz (*alpha*) band whereas dorsal stream areas prefer tracking > 14 Hz (*beta*) dynamics in our case. This finding may point towards a frequency-specialization of tracking different temporal dynamics. A “spectral specialization” of brain areas and networks has further been concluded from recent findings from MEG human resting state activity at the whole brain level (Groppe et al., 2013, Keitel & Gross, 2016) and resonance to magnetic pulse stimulation (Rosanova et al., 2009). Consequentially, cortices with a characteristic spectral fingerprint might preferentially respond to external stimulation that makes pace in their resonant frequency range. In case of the visual system, evidence suggests that the occipito-parietal alpha-rhythm can be entrained by alpha-rhythmic visual stimulation (Gulbinaite, Roozendaal, & VanRullen, 2019; Haegens & Zion Golumbic, 2018; Notbohm & Herrmann, 2016; Spaak, de Lange, & Jensen, 2014; Thut, Schyns, & Gross, 2011).

In our case however, visual cortices generally responded to all three quasi-rhythmic stimulation modes, thus challenging a special role of alpha stimulation (also see Keitel, Benwell, Thut, & Gross, 2018). Areas resonating within certain frequency bands, such as parietal cortex within the alpha range (8-13 Hz) and motor cortex within the beta range (around 20 Hz), did not show tracking of the respective stimulus temporal dynamics (Fig. 5). Further note that frontal areas failed to track the temporal dynamics as shown in two analyses: when pooling coupling measures for attended and non-attended stimuli (Fig 5a) and in a more sensitive control analysis (see Methods section for details) using only coupling measures for attended stimuli (Supplementary Fig. 4). This is in contrast with results from previous studies, which suggest that some frontal visual areas like the frontal eye fields (FEF) may be functionally situated at an intermediate or even early processing stage of the visual hierarchy (Bastos et al., 2015). We emphasize however that our findings do not exclude the possibility that frontal areas, such as FEF, have a more indirect role in processing quasi-rhythmic visual input, either by aggregating information about temporal dynamics from lower visual areas or by exerting top-down control, as an example, by modulating tracking by attention. Also note that the involvement of these areas may depend on the immediate task-relevance of the temporal structure of the stimuli: Our stimuli enable a tracking of visual processing at the stimulus location but they are not temporally predictive of target presentations. It is possible that using them as predictors, by e.g. presenting stimuli at a certain phase (see Mathewson et al., 2012; Sokoliuk & VanRullen, 2016), frontal areas exert quasi-rhythmic activity that reflects top-down predictions about imminent target presentation times.

Finally, we also found a few ipsilateral cortices tracking temporal dynamics, mostly in the *theta* condition (Supplementary Fig. 1). Being outside the longitudinal fissure, these effects cannot be explained by a spread due to the limited spatial resolution of the beamformer source reconstruction or by errors in segmenting ventral cortex by using T1-weighted MR anatomical scans. Previous investigations using quasi-rhythmic visual stimulation in the *theta* frequency band (Biau, Morís Fernández, Holle, Avila, & Soto-Faraco, 2016; Hauswald, Lithari, Collignon, Leonardelli, & Weisz, 2018; Park, Kayser, Thut, & Gross, 2016) did not differentiate between ipsi- and contralateral cortical processing due to non-lateralised visual stimulation. Thus, further investigations are required to better elucidate the role of these ipsilateral areas.

Taken together these findings suggest that the temporal structure of visual input is processed similarly in primary and early extra-striate visual areas but differences arise between stimulation frequency ranges at later stages. The observed differences might reflect different processing networks corresponding to different ecologically relevant frequency bands. Our data support this conclusion, especially considering that maximum tracking occurred for theta band stimulation, a range that has been found to underlie and support cognitive function in everyday life situations, such as understanding speech (Park et al., 2016; Marinato & Baldauf, 2019), interpreting hand gestures (Biau et al., 2016) and monitoring peripheral vision during locomotion.

Note however that a potential alternative explanation for the far spread of theta tracking is its relative strength in sensor space. Given that we used a common spatial filter for all experimental conditions, strong theta tracking may have contributed to its spatial spread in source reconstructions. This point could be addressed in follow-up work by factoring in the perceptual relevance and behavioral consequences of each tracking response (Keitel, Gross, & Kayser, 2018) or, as pointed out by a reviewer, applying leakage correction (such as orthogonalisation) in source reconstruction (Colclough, Brookes, Smith, & Woolrich, 2015). Moreover, it has to be noted that coherence at a given frequency, while being highly sensitive to phase locking, also depends on signal to noise ratio (SNR), mainly because of the interference of spontaneous activity at that frequency (Srinivasan, Russell, Edelman, & Tononi, 1999). While it is difficult to estimate SNR for real data, this effect could play a role in the difference of tracking extent we observe in different flickering conditions, in particular considering that, most probably, SNR is greater for the theta band (because of 1/f noise) and for the alpha band (because of spontaneous activity).

### 4.2 Attention modulates coherence in all quasi-rhythmic stimulation conditions

We compared the cortical distribution and magnitude of the attentional modulation of quasi-rhythmic brain-stimulus coupling between theta, alpha and beta stimulation conditions. In each condition we observed enhanced cortical tracking of attended stimuli in at least one visual area. Primary visual cortex V1 was the only cortical area that showed systematic gain effects with comparable magnitude in all three conditions (as well as in the SSR condition – see following section). Once more, this finding highlights the role of V1 as the first cortical locus prioritizing the processing of task-relevant visual information (Somers, Dale, Seiffert, & Tootell, 1999, Martínez et al., 1999, Buffalo, Fries, Landman, Liang, & Desimone, 2010). It further corroborates V1’s role as an important hub in monitoring and controlling the processing of continuous dynamic visual input (Gandhi, Heeger, & Boynton, 1999), and given the nature of our stimuli, extends this role to include quasi-rhythmic dynamics within distinct frequency bands below 20 Hz. The specific design of our stimuli, essentially a type of contrast-modulated Gabor patches, and the here implemented change-detection task may have further promoted the general involvement of V1 in modulating visual input (Glickfeld, Histed, & Maunsell, 2013). Behavioral performance data and an alpha-lateralization control analysis confirmed that participants where indeed allocating and maintaining their spatial attention as instructed throughout the experiment.

Compared with alpha and beta conditions, we observed the cortically most widespread attention effect during theta stimulation. In addition to V1, attention enhanced the cortical representation of stimulus dynamics in all early visual areas V2-V4 and in close neighbors in posterior cingulate cortex (areas POS1 and DVT) and MT+ Complex (V4t). Further effects were found along the ventral stream with an emphasis on the fusiform cortex (FFC) and adjacent areas (VMV1, TE2p). Attention effects in the dorsal stream were restricted to area IPS1 that borders on the Dorsal Visual Transition area (DVT), which in turn is a direct neighbor of V2. Of particular interest is the involvement of the FFC and other areas of the ventral stream. One function of the ventral stream is to successively integrate low level visual features to high-level visual objects (Milner & Goodale, 2006), with regions such as FFC showing a preference for processing object classes (e.g. faces in its fusiform face area, FFA). Several experimental studies (Kanwisher, 2010, Coggan, Baker, & Andrews, 2019, Baldauf & Desimone, 2014; DeVries & Baldauf, 2019) have shown that parts of the FFC follow relatively slow fluctuations (~2 Hz) during rhythmic face presentation and that this response was enhanced when participants attended to the face presentation. Our results suggest that the FFC (and its neighbors) allow tracking of temporally slow fluctuations in visual objects more generally. In this case, the occurrence of attentional modulation during theta, but not during faster alpha and beta stimulation could have been a consequence of an increasing sluggishness of higher order ventral areas to follow stimulus modulations or, put differently, longer temporal integration windows for visual input (Gauthier, Eger, Hesselmann, Giraud, & Kleinschmidt, 2012). Note that theta stimulation also produces the spatially most extensive propagation of cortical tracking as described in the previous section. Observing the most widespread distribution of attentional modulation for theta could thus be a consequence of this propagation effect. Taking into account that theta produces the strongest brain-stimulus coupling in general the possibility remains that reported differences are due to signal-to-noise issues rather than a preference of the visual system for theta band fluctuations in visual input (Srinivasan, Russell, Edelman, & Tononi, 1999).

Regarding brain-stimulus coupling in the alpha condition, attentional modulation was confined to V1 despite its relatively widespread cortical tracking. Although we cannot rule out a functional difference for stimulation in this frequency range, a likely explanation is that measuring cortical tracking in this condition suffers from cross-talk with ongoing alpha-rhythms. Visual cortex has long been known for exhibiting strong intrinsic rhythmic activity in the ~10 Hz range (Palva & Palva, 2007, Keitel & Gross 2016). Most, if not all, intrinsic alpha-rhythmic activity generated in visual cortex occurs at random phases relative to the stimulation. Occasionally, intrinsic and stimulus-driven rhythms will align (or oppose) their phase by chance and produce a biased estimate of our phase-based measure of brain-stimulus coupling. Because intrinsic alpha activity shows a similar but reversed lateralisation during focused visuo-spatial attention – alpha decreases (i.e. is suppressed) contralaterally to the attended location and vice versa (Capilla, Schoffelen, Paterson, Thut, & Gross, 2014; Keitel, Benwell, Thut, & Gross, 2018; Kelly, Lalor, Reilly, & Foxe, 2006; Thut, Nietzel, Brandt, & Pascual-Leone, 2006) – it is possible that contralateral gain and alpha suppression effects cancel each other out in some areas. Not suffering from this caveat, the attentional modulation of beta-band tracking was more far-spread although not comparable with the spatial distribution of theta gain effects: In addition to V1, early visual cortex V2, dorsal stream area V3A and ventral stream area V8 showed gain effects.

What are the neurophysiological principles underlying the attentional gain effect and its behavioural consequences? The latter is out with the scope of the present study because participants were instructed only to respond to target events on the attended stimulus. This precludes further analysis that would allow us to characterize effects as an increase in either accuracy or response bias (Benwell et al., 2017). Follow-up studies could adapt our paradigm to a Posner-type situation in which validly and invalidly cued targets and different contrast levels enable a more detailed analysis of behavioural performance, As for the neurophysiological underpinnings of increased tracking for attended stimulation, it seems straightforward to turn to the classical sensory gain perspective: attended stimuli win a competition for neural processing resources awarding them a prioritized cortical representation (Desimone & Duncan, 1995; Kastner & Ungerleider, 2001). However, sensory gain is almost exclusively based on studies that use static and/or transient stimulation. Little is known about how these principles apply during sustained attention to continuous (quasi-rhythmic) visual stimulation. In the simplest case a constant gain is applied to attended visual input over time as suggested by current “static” computational models of attention (Boynton, 2009; Reynolds & Heeger, 2009) although cortical signals might be more complicated: Recent evidence strongly suggests that attentional allocation itself is dynamic, if not inherently rhythmic (Helfrich et al., 2018; Senoussi, Moreland, Busch, & Dugué, 2019) and a dynamic sensory gain or normalization theory of attention has not yet been formulated (for a starting point see Montijn et al. 2012).

Alternatively, it may help regarding the attention effect as an increase in fidelity, i.e. the faithfulness with which the neuronal populations follow the dynamics of their preferred feature (Chennu, Craston, Wyble, & Bowman, 2009). This could be intimately linked to a release from a strong intrinsic regime as we have recently argued (Keitel et al. 2019): Also in line with the present data alpha, as an internal synchronizer, is suppressed in cortical regions that process attended locations. This release from an internal pacemaker allows affected neuronal populations to couple more strongly to stimulus dynamics. Further evidence for this perspective comes from observations that stimulus presentation itself and the allocation of attention both reduce neural variability, a measure thought to reflect an increase in cortical signal-to-noise ratio (Arazi, Censor, & Dinstein, 2017; Arazi, Yeshurun, & Dinstein, 2019; Deco & Hugues, 2012). Interestingly, in turn, changes in neural variability seem to be tied most closely to fluctuations in the power of alpha rhythms (Daniel, Meindertsma, Arazi, Donner, & Dinstein, 2019).

### 4.3 Attention modulates coherence with strictly rhythmic stimulation in primary and early visual areas

The present experiment featured an additional strictly-periodic stimulation condition. The left stimulus elicited a 10 Hz steady-state response (SSR), predominantly in the contralateral (right) visual cortex. The right visual stimulus elicited a 12 Hz SSR, largely generated in left visual cortices due to the spatiotopic organization of the visual system. As previously reported a variety of areas, from primary visual up into dorsal and ventral streams contributed to the macroscopic SSRs (Di Russo et al., 2007; Fawcett, Barnes, Hillebrand, & Singh, 2004; Pastor, Artieda, Arbizu, Valencia, & Masdeu, 2003). Our visuo-spatial attention manipulation replicated previous reports of attentional gain effects on an inter-trial phase-consistency based measures of SSRs (Gulbinaite, Roozendaal, & VanRullen, 2019; Joon Kim, Grabowecky, Paller, Muthu, & Suzuki, 2007; Kashiwase, Matsumiya, Kuriki, & Shioiri, 2012; Porcu, Keitel, & Müller, 2013) that is mathematically equivalent to the here employed phase cross-coherence measure of brain-stimulus coupling.

Our whole-brain analysis, supported by a fine-grained cortical parcellation (Glasser et al. 2016), allowed us to investigate the cortical topography of attentional modulation with unprecedented spatial detail. The results support an attentional modulation of SSRs as early as V1 (Lauritzen et al. 2010, Palomares et al. 2012, Keil et al. 2012) and substantiate attentional gain effects in subsequent early visual cortices V2, V3 and V4, as well as in dorsal stream area V6 and ventral stream area V8. In addition to V1 and V4, Lauritzen et al. (2010) reported SSR attentional gain in the middle temporal area (hMT+ in their notation, likely a region in the MT+ complex of Glasser et al. 2016) and in the intraparietal sulcus (IPS in their notation, possibly IPS1 in Glasser et al. 2016), which we did not find. However, Lauritzen et al. (2010) pre-defined ROIs functionally (hMT+) or anatomically (IPS). Therefore, effects might be explained by spill-overs from adjacent areas V6 (in case of IPS) and V4 (hMT+, via V4t) that were not part of their analysis but might have been included due to spatial uncertainty or co-activation.

Two additional areas were found to increase stimulus-coupling with attention: 7Am in the Superior Parietal Cortex (SPC) and p24 in the Anterior Cingulate Cortex (ACC). Both regions, the ACC (e.g. Silton et al., 2010) and the SPC (e.g. Sereno, 2001) have been implicated in the allocation of attention. However, ascribing such a function to respective sub-units here remains speculative because p24 and 7Am are relatively isolated and do not show as a locus of attentional modulation in the quasi-rhythmic stimulation conditions.

Finally, the detailed source reconstruction allows for re-visiting attentional effects on SSRs recorded with EEG and analyzed on the scalp-level. Using a similar paradigm, Keitel et al. (2017, also see Keitel et al. 2019) have described counter-intuitive topographical patterns of attention effects on 10- and 12-Hz SSRs. Although SSRs themselves were maximal at EEG recording sites contralateral to the stimulus location, attention effects were not strictly co-localized with these maxima but seemed to occur non-lateralized or even ipsilateral based on their projection to the scalp. However, inferring cortical loci from scalp topographies can be deceiving. The present source-level analysis demonstrates that the attentional modulation is predominantly contralateral as expected. Previously described topographical mismatches could stem from the fact that not all of the areas that track stimulus dynamics also show attentional modulation. Moreover, some of the areas that are subject to attentional modulation lie within (V1) or close to the Calcarine Fissure (V2, V4). These anatomical constraints make a strongly (contra-) lateralized scalp projection in EEG measurements less likely.

### 4.4 Coherence analysis of quasi-rhythmic stimuli disentangle contribution in the same frequency band

Our methodological approach allowed us to spectrally and spatially separate the cortical tracking of concurrently presented stimuli oscillating in the same frequency band. Brain responses were isolated and specific to the actually contralateral presented peripheral stimulus, as demonstrated in Fig. 4a. This is consistent with the spatiotopic organization of early visual cortices, also considering that residual ipsilateral brain-stimulus coupling localized to the longitudinal fissure is most likely due to a contamination effect from the beamformer solution (Van Veen, Van Drongelen, Yuchtman, & Suzuki, 1997). This last aspect points to a limitation of the method in differentiating lateralized contributions, especially for primary and early visual areas that fall within the longitudinal fissure. Another limitation lies in the fact that classical delay and phase analysis is complicated by the continuous change in frequency of the presented stimulus. Finally, given our data alone we cannot yet uniquely attribute the effect that theta stimulation produced the strongest and most wide-spread tracking to the specific frequency range per se. The magnitude of brain-stimulus coupling in the quasi-rhythmic regime might also be a function of the stimulus bandwidth, which was 3 Hz for *theta* and increased up to 6 Hz for *beta* stimulation. Animal single cell studies however suggest that the primary visual cortex has a propensity to code natural broad-band visual dynamics in LFPs with fluctuations < 12 Hz (Mazzoni, Brunel, Cavallari, Logothetis, & Panzeri, 2011), thus possibly explaining the gradient that we observe without the need to assume a specific role for stimulus bandwidth.

Apart from these limitations, our approach provides a means for investigating brain responses to temporally dynamic stimuli. The classical approach of strictly rhythmic frequency tagging requires the frequencies of presented stimuli to be well separated, unless recently developed spatial filtering algorithms are applied that allow for a closer spacing of stimulation frequencies (Cohen & Gulbinaite, 2017; Nikulin, Nolte, & Curio, 2011).

The present approach, based on expressing brain-stimulus coupling in terms of a phase cross-coherence measure can be useful to investigate brain responses to more complex and naturalistic stimuli while preventing perceptual differences between them. Moreover, covering frequency bands instead of a single frequency, our method allows for investigating entrainment phenomena by tailoring the stimulation to neurophysiologically well-defined frequency ranges such as alpha.

## 5. Conclusion

By investigating phase synchronization in MEG-recorded and source-projected cortical activity, we found that many occipital, parietal and temporal areas of the brain tracks the temporal structure of quasi-rhythmic stimuli oscillating in classical theta, alpha and beta frequency bands. Moreover, we found that focused attention enhances the coupling between stimulus contrast modulation and neural activity. Using a state-of-the-art parcellation of the human cortex (Glasser et al. 2016), we were able to provide a spatially detailed characterization of which cortical regions are involved in the tracking of strictly- and quasi-rhythmic stimulation, and their modulation by attention. Crucially, the extent and the magnitude of the synchronization showed differences between stimulation frequencies – being strongest and most widespread during theta stimulation. This may point towards frequency specialized networks for the processing of visual dynamics. Additionally, we developed and validated a method allowing for tracking the contribution of concurrently and peripherally presented stimuli, fluctuating within the same frequency band. This type of stimulation has the advantages of being perceptually equivalent and resembling ecologically relevant visual input, such as speech reading, hand gestures and peripheral vision in locomotion, in a controlled manner.

## Data Availability Statement

Data will be made available upon reasonable request. Source code used for the analysis is publicly available on Open Science Framework (<u>DOI 10.17605/OSF.IO/7E9C2</u>) under the GNU General Public License (GPL) 3.0

## Acknowledgments

We would like to thank the research groups of Uri Hasson. Angelika Lingnau, Olivier Collignon and Scott Fairhall for sharing previously acquired anatomical MRI scans for some of our experimental subjects. DT was funded in part by the Erasmus+ program of the European Union. CK and JG were funded by a Wellcome Trust Senior Investigator Grant awarded to JG (#098433). CK received support from Wellcome Trust ISSF Secondment (204820/Z/16/Z) and BBSRC Flexible Talent Mobility funding (BB/R506576/1) awarded by the University of Glasgow.

## Supplementary Figures

**Supplementary Fig 1:**
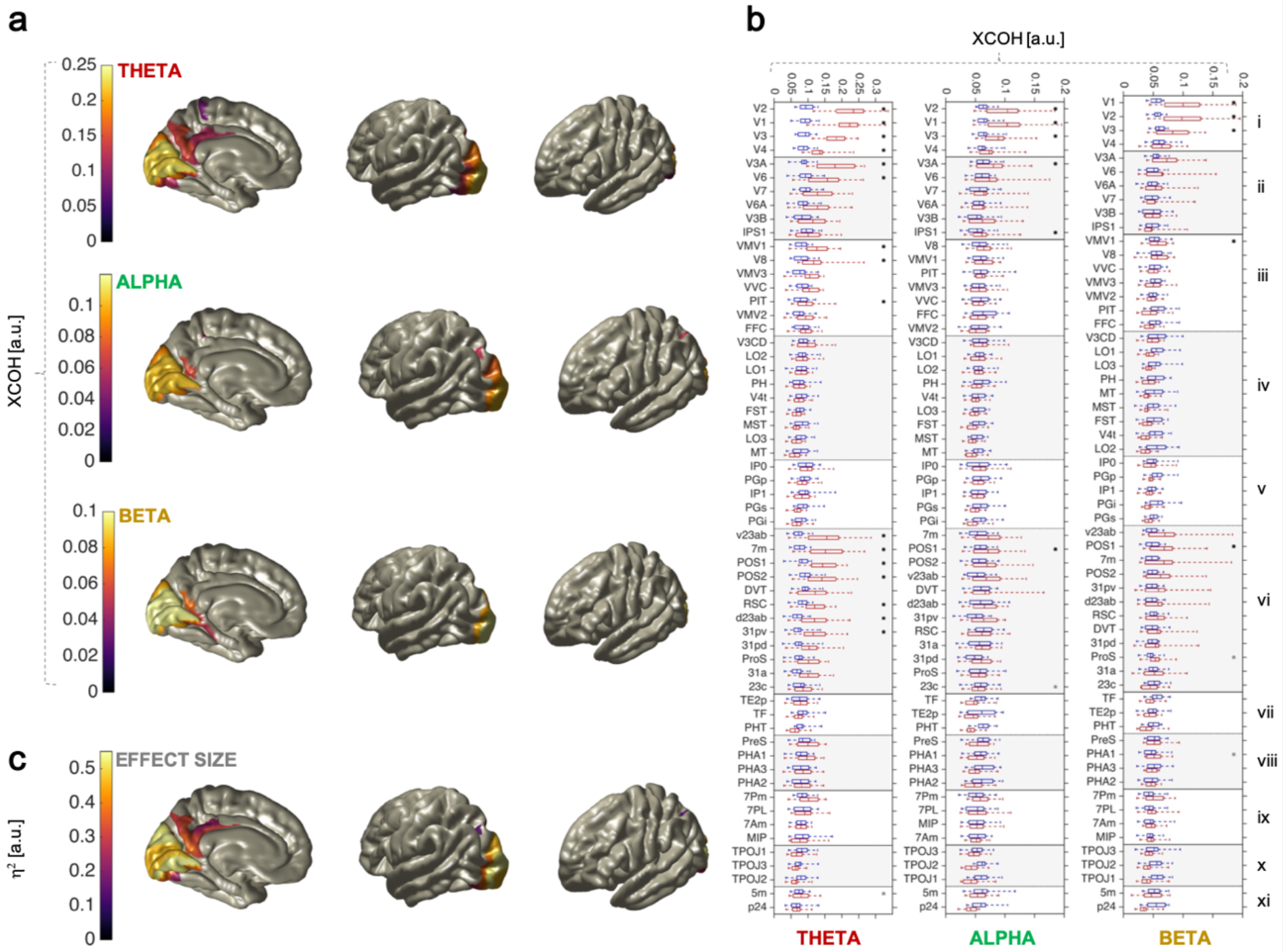
Ipsilateral cortical tracking of underlying quasi-rhythmic temporal dynamics. In (a) grand-average ROI-specific cross-coherence (XCOH) with ipsilateral stimulus 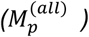 is reported for all quasi-rhythmic stimulation conditions; each cortical ROI is colored according to its correspondent value; data collapsed across attention conditions; ROIs are masked by significance (Bonferroni corrected p-value < 0.01, 180 comparisons) from a one-sided non-parametric dependent samples t-test comparing Fisher stabilized XCOH with actual 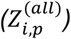 and surrogate 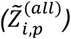 contrast modulations. Asterisks on boxplots in (b) indicates areas significantly tracking the contrast underlying dynamics according to the above mentioned statistical comparison; boxplots show XCOH with actual (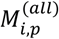 in red) and surrogate contrast (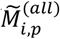 in blue) for each ROI in Table 1; roman numerals identify ROI district, as per Table 2; black asterisks indicate that the magnitude of XCOH further depends on the stimulation frequency, as determined by a one-way repeated-measures ANOVA on Fisher stabilized XCOH differences 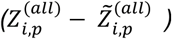 between actual and surrogate data. Corresponding effect sizes are reported in (c) only including ROIs whose XCOH depended on the stimulation frequency. Stimulus frequency has largest effects on XCOH in early visual cortices.

**Supplementary Fig 2:**
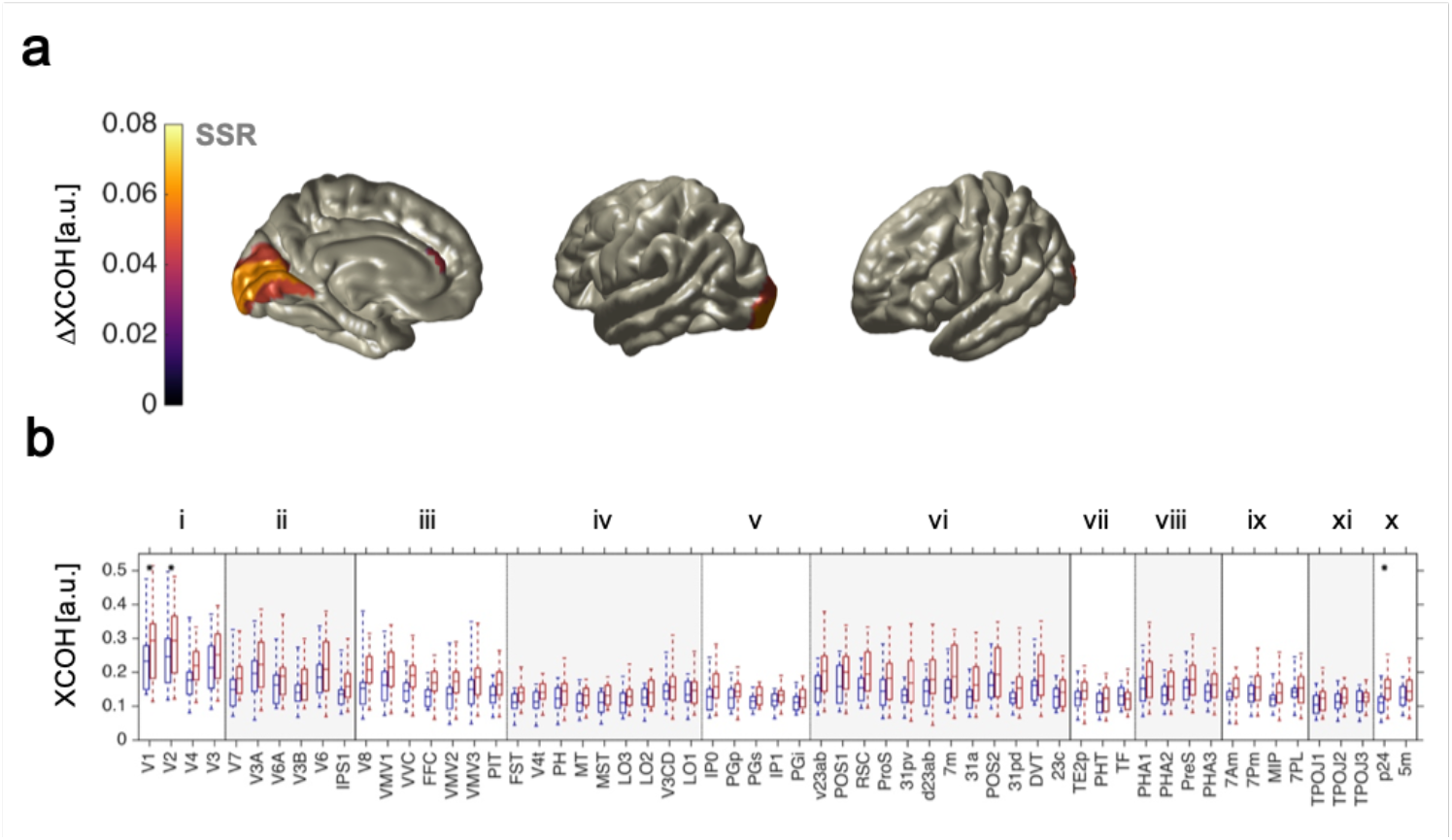
Ipsilateral attentional modulation of Steady State Responses. In (a) the difference 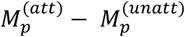 of grand-average ROI-specific cross-coherence (XCOH) with a ipsilateral attended and unattended stimulus is shown; each ROI on the cortex is colored according to the difference in XCOH (= ΔXCOH); only significant differences are displayed (Bonferroni corrected p-value < 0.05, 180 comparisons) according to a one-sided non-parametric dependent-samples t-test comparing Fisher stabilized XCOH in the attended 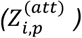 and unattended 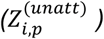 condition. Asterisks on boxplots in (b) indicates areas where the XCOH with the contrast modulation is significantly enhanced as per the above statistical comparison; boxplots show XCOH in the attended (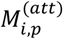 in red) and unattended (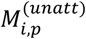 in blue) condition for each ROI in Table 1; roman numerals identify ROI districts, as from Table 2.

**Supplementary Fig 3:**
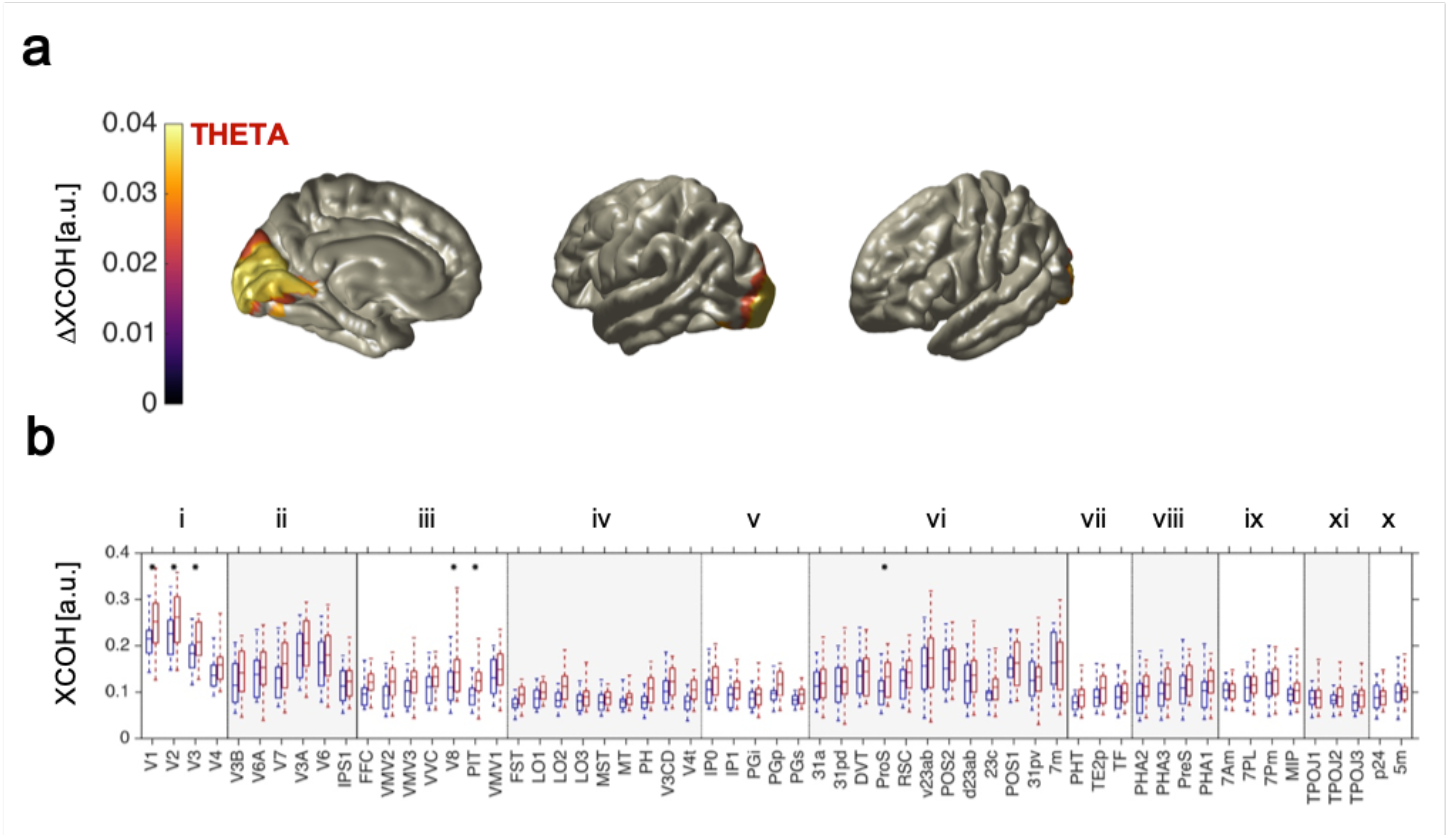
Ipsilateral attentional modulation in quasi-rhythmic conditions. In (a) the difference 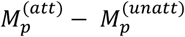 of grand-average ROI-specific cross-coherence (XCOH) with ipsilateral attended and unattended stimulus is shown; each ROI on the cortex is colored according to the difference in XCOH (= ΔXCOH); only significant differences are displayed (Bonferroni corrected p-value < 0.05, 21 comparisons), according to a one-sided non-parametric dependent-samples t-test comparing Fisher stabilized XCOH in the attended 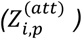 and unattended 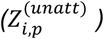 condition. Asterisks on boxplots in (b) indicates areas where the XCOH with contrast modulation is significantly enhanced as per the above statistical comparison; boxplots show XCOH in the the attended (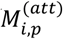 in red) and unattended (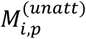 in blue) condition for each ROI in Table 1; roman numerals identify ROI districts, as per Table 2.

**Supplementary Fig 4:**
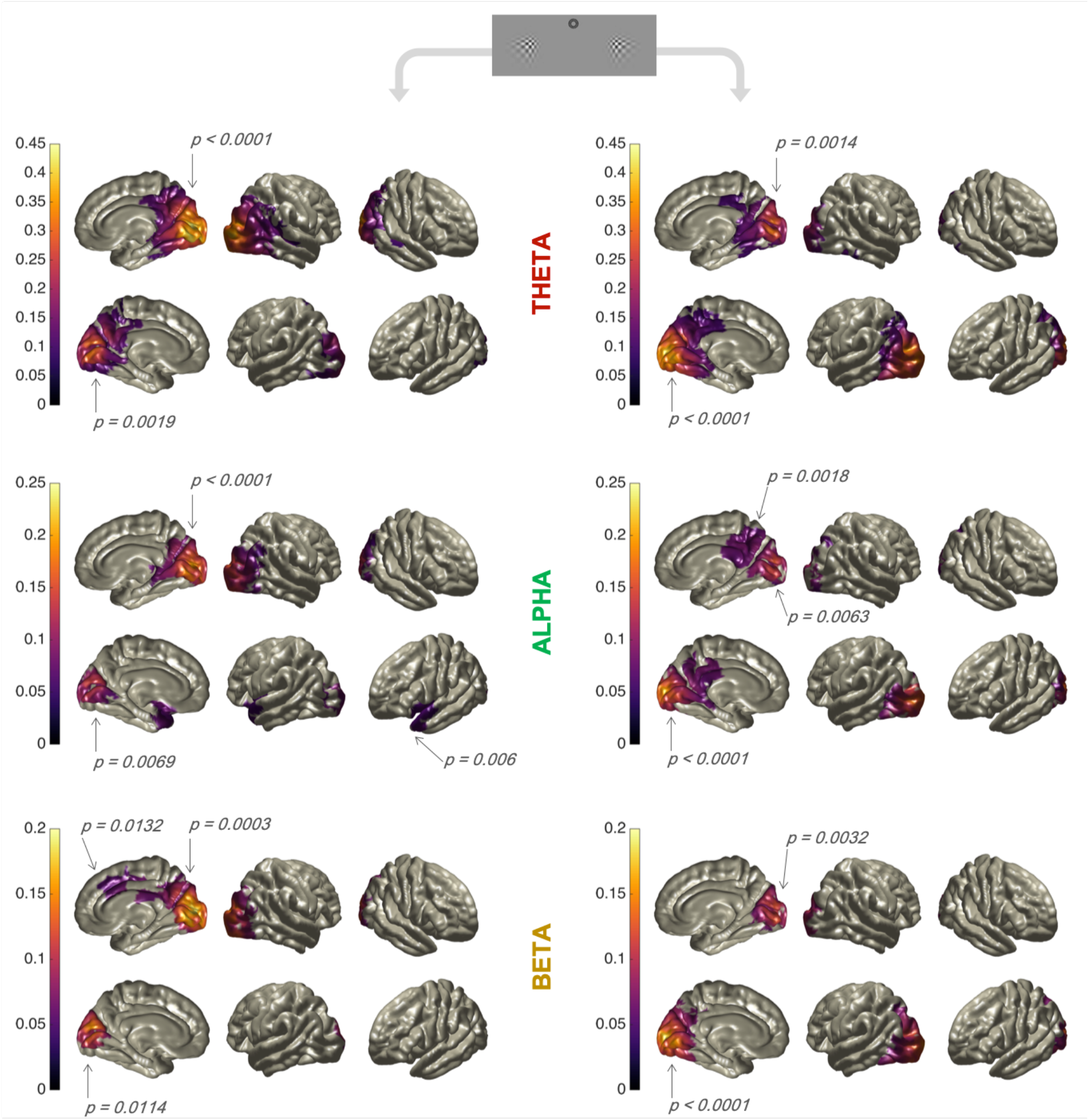
Dipole-based analysis of cortical tracking of underlying quasi-rhythmic temporal dynamics. Cross coherence with left/right contrast modulation was computed using only trials in which the respective driving stimulus was attended. Results were masked according to a non-parametric one-tailed dependent samples t-test, with cluster-based correction, comparing Fisher stabilized XCOH with actual 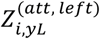 and 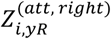 and corresponding surrogate (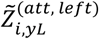 and 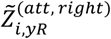) contrast modulations. Only clusters with p < 0.01 (or slightly above this threshold) are shown. Notably, all other clusters have p > 0.55. Arrows indicate p-value of respective clusters. Right/left hemisphere corresponds to contra/ipsilateral results, respectively.

**Supplementary Fig 5:**
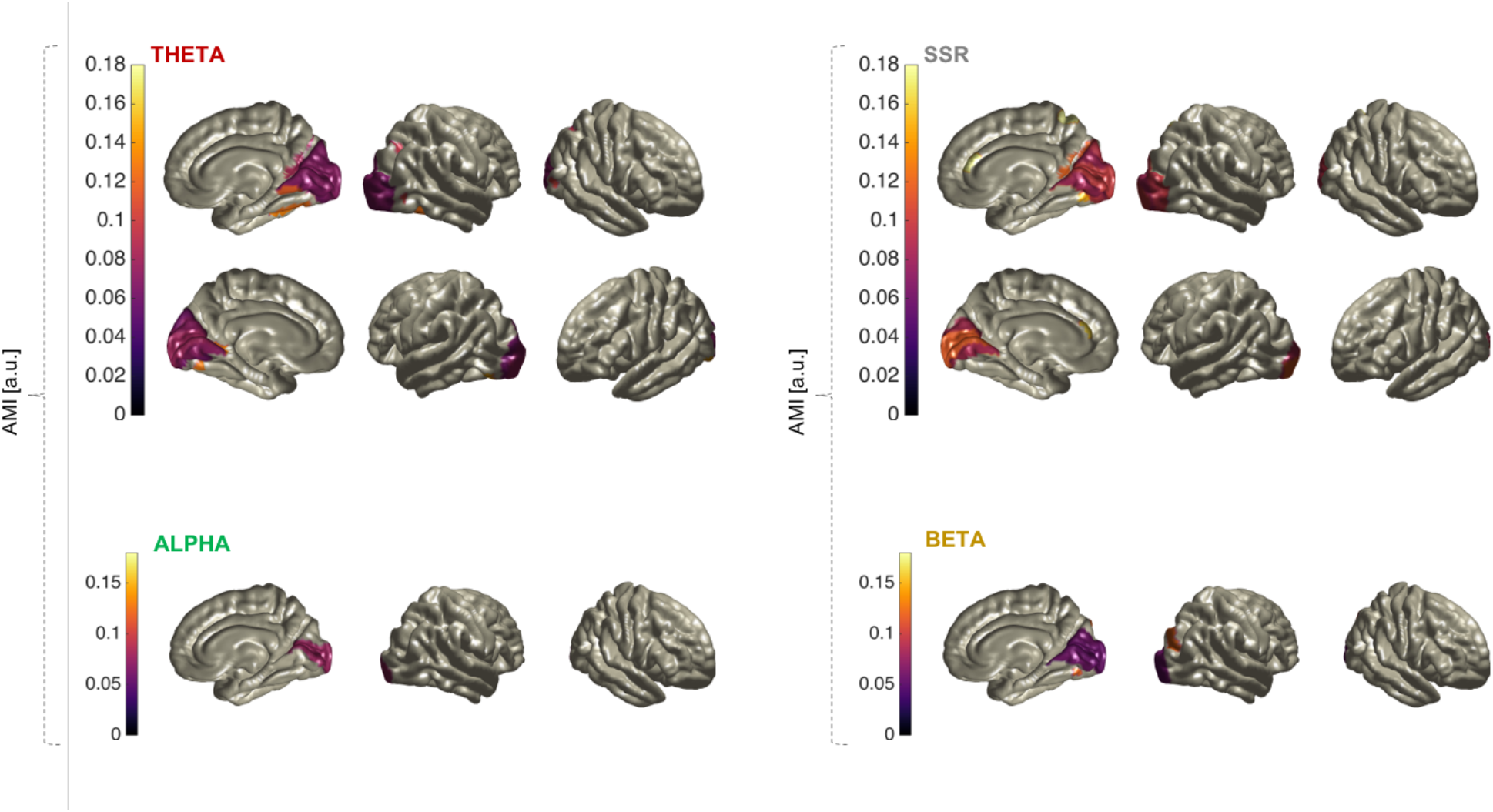
Attentional modulation index 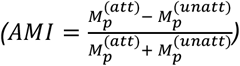 for the ROIs showing a significant difference of coherence when comparing attended/unattended trials in all the flickering conditions. AMI has been computed for ROIs showing significant attentional modulation in the Fisher-stabilized cross coherence analysis (see Fig. 5 and Fig 6 for details). Right/left hemisphere corresponds to contra/ipsilateral results, respectively.

